# Disulfiram reduces atherosclerosis and enhances efferocytosis, autophagy, and atheroprotective gut microbiota in hyperlipidemic mice

**DOI:** 10.1101/2023.10.17.562757

**Authors:** C. Alicia Traughber, Kara Timinski, Ashutosh Prince, Nilam Bhandari, Kalash Neupane, Mariam R Khan, Esther Opoku, Emmanuel Opoku, Gregory Brubaker, Kanuri Nageshwar, Elif G. Ertugral, Prabha Naggareddy, Chandrasekhar R. Kothapalli, Jonathan D. Smith, Kailash Gulshan

## Abstract

Pyroptosis executor Gasdermin (GsdmD) promotes atherosclerosis in mice and humans. Disulfiram (DSF) was recently shown to potently inhibit GsdmD, but the in-vivo efficacy and mechanism of DSF’s anti-atherosclerotic activity is yet to be explored. We used human/mouse macrophages and a hyperlipidemic mouse model of atherosclerosis to determine DSF anti-atherosclerotic efficacy and mechanism. DSF-fed hyperlipidemic apoE^-/-^ mice showed significantly reduced IL-1β release upon in-vivo Nlrp3 inflammasome assembly and showed smaller atherosclerotic lesions (∼27% and 29% reduction in males and females, respectively). The necrotic core area was also smaller (∼50% and 46% reduction in DSF-fed males and females, respectively). DSF induced autophagy in macrophages, hepatocytes/liver, and in atherosclerotic plaques. DSF modulated other atheroprotective pathways such as efferocytosis, phagocytosis, and gut microbiota. DSF-treated macrophages showed enhanced phagocytosis/efferocytosis, with a mechanism being a marked increase in cell-surface expression of efferocytic receptor MerTK. Atomic-force microscopy analysis revealed altered biophysical membrane properties of DSF treated macrophages, showing increased ordered-state of the plasma membrane and increased adhesion strength. Furthermore, the 16sRNA sequencing of DSF-fed hyperlipidemic mice showed highly significant enrichment in atheroprotective gut microbiota *Akkermansia* and a reduction in atherogenic *Romboutsia* species. Taken together, our data shows that DSF can simultaneously modulate multiple atheroprotective pathways, and thus may serve as novel adjuvant therapeutic to treat atherosclerosis.

## Introduction

Atherosclerosis, a sterile and chronic inflammatory disease, is the major cause of cardiovascular disease (CVD) related mortalities^1^. The complex nature of atherosclerosis is highlighted by the involvement of multiple pathways such as cholesterol efflux, NLRP3/AIM2 inflammasomes, neutrophils extracellular traps (NETosis), autophagy, efferocytosis, and gut microbiota^1–5^. Cumulative defects or dysregulation of these pathways over time promotes progression of atherosclerosis. Thus, tackling CVD with cholesterol-lowering therapeutics, such as statins, alone may not be enough to reverse the trend of increasing prevalence of CVD. New CVD therapeutics are needed to target multiple pathways simultaneously, and that can be used in conjunction with cholesterol-lowering therapies. Human trials showed that IL-1β might serve as one of the targets for adjuvant therapy for CVD^6–8^. IL-1β acts as a local vascular and as a systemic contributor in pathogenesis of CVD^1,7–9^. The generation of biologically active IL-1β in atherosclerotic plaque area depends on activation of the Nlrp3 inflammasome^10–12^. More recently the AIM2 inflammasome was also identified as an age-related risk factor for CVD^13^, pointing towards a sustained role of inflammasomes in exacerbating atherosclerosis in humans. Activation of inflammasomes leads to cleavage of pore-forming protein Gasdermin D (GsdmD) in plaques, allowing local as well as systemic release of proinflammatory cytokines, which further exacerbates disease via recruitment of new immune cells to plaque areas^14–16^. Recent studies highlight the atherogenic role of GsdmD^13–17^, with GsdmD mRNA being up regulated in peripheral blood monocytes of patients with coronary artery disease, and the presence of cleaved N-terminal (NT)-GsdmD fragment in mouse and human atherosclerotic plaques^14–16^, but the mechanism of atherogenic activity of GsdmD is still not fully resolved. GsdmD can be cleaved via capase-1/Nlrp3 dependent canonical inflammasome pathway, or via a caspase-11 (in mice) or caspase-4/5 (in humans) dependent non-canonical inflammasome pathway. The NT-GsdmD fragment generated via non-canonical pathway can in-turn activate the canonical inflammasome in a positive feedback loop^18^. GsdmD thus serves as a common executor of both canonical and non-canonical pyroptosis^19–22^.

Targeting GsdmD seems to be a better option than anti-IL-1β, as GsdmD mediates highly inflammatory pyroptosis, which is upstream of IL-1β release. This leads to a large increase in danger-associated molecular patterns (DAMPs) molecules that can further amplify inflammation in atherosclerotic plaques. Blocking GsdmD can reduce the levels of not only IL-1β, but also of IL-18 and other DAMPs. Furthermore, studies have shown role of GsdmD in formation of neutrophils extracellular traps (NETs)^2,23,24^, thus blocking GsdmD can also reduce atherogenic NETosis. In contrast to GsdmD/IL-1β atherogenic role, autophagy and efferocytosis serve as major atheroprotective pathways. Defective autophagy was observed in advanced human and mouse atherosclerotic lesions, while up regulation of autophagy was shown to protect against CVD^3,25–28^. Efficient efferocytosis of damaged cholesterol-laden foam cells is one of the strategies used by the host immune system to promote regression of atherosclerotic plaques and prevent CVD^4,29–32^. In addition to intracellular atheroprotective pathways, the gut microbiota and associated metabolites can modulate the host’s risk of developing CVD, independent of traditional risk factors^33^. One striking example is the gut microbiota mediated conversion of dietary choline into atherogenic trimethyl amine oxide (TMAO), which promotes atherosclerosis in humans^5,34–36^. Thus, the next generation of statin-adjuvant CVD therapeutics are expected to target inflammasomes, GsdmD, NETosis autophagy, efferocytosis, and gut microbiota.

The FDA-approved drug Disulfiram (DSF) was shown to potently inhibit GsdmD pore formation and block formation of NETs^36, 37^, but its anti-atherosclerotic effect and mechanism are not clear. Here, we report that DSF reduces atherosclerotic burden and necrotic core area in plaques in a hyperlipidemic mouse model of atherosclerosis. DSF causes plasma membrane remodeling and induces anti-atherosclerotic pathways such as autophagy and efferocytosis. Furthermore, DSF promotes enrichment of atheroprotective gut microbiota and reduction in atherogenic bacterial species in hyperlipidemic mouse model of atherosclerosis.

## Methods

### Cell culture

Cell culture and treatment conditions for RAW-ASC, RAW-ASC-GsdmD^-/-^, THP-1, Jurkat, and HepG2 cells are described in *supplementary material and methods* section. Bone marrow derived-macrophages (BMDMs) were isolated and cultured as described earlier^14^.

### In vitro and In-vivo inflammasome assembly

In vivo Nlrp3 inflammasome assembly, in-vitro Nlrp3 inflammasome assembly, and IL-1β release assays were performed as described earlier^14^. Detailed methods are described in *supplementary material and methods* section.

### Viability Assay

Viability of cells was measured by Live/dead fixable blue stain kit, with detailed method in *supplementary material and methods* section.

### Cytotoxicity Assay

Toxicity of DSF on cells was measured by the CyQuant LDH Cytotoxicity Assay kit (ThermoFisher), with detailed method in supplementary material and methods section.

### Mice and Diets

Animal experiments were performed in accordance with approved protocols from the Cleveland State University and the Cleveland Clinic Institutional Animal Care and Use Committees. The C57BL6J-WT mice were purchased from the Jackson Laboratory and the C57BL/6J-GsdmD^−/−^mice were generated earlier^37^ and kindly provided by Dr. Russell Vance (UC, Berkeley). C57BL/6J-apoE^-/-^ mice were purchased from Jackson Laboratories and bred in house. Mice were maintained in a temperature-controlled facility with the standard 12-hour light/dark cycle.

### Atherosclerosis studies

To promote hyperlipidemia, mice were weaned onto a Western Type Diet (WTD) containing 42% calories from fat as described by others^38–40^. Disulfiram (Sigma; #PHR 1690) was milled into WTD at 100mg/kg/day by Envigo, as described previously^41^.

### Blood collection and analyses

EDTA-anticoagulated blood samples were collected via tail vein bleed every three weeks post diet start and by retro-orbitally at endpoint. Plasma was isolated by centrifugation at 15,000 rpm for 5 min and stored at −80°C until assayed. Total cholesterol was measured by using Stan Bio Total cholesterol reagent (#1010-225), following manufacturer’s instructions. Endpoint whole blood was diluted using 3% BSA prepared in PBS. Blood was analyzed in a complete blood count panel on the Siemens Advia 120 Hematology Analyzer at the Cleveland Clinic Laboratory Diagnostics Core. Endpoint plasma samples were subjected to FPLC, in which 100 μl of pooled plasma from respective groups were separated on a Superose 6 column 10/30 (Cytiva).

### Atherosclerotic Lesion and Necrotic Core Microscopy and Quantification

Mice were sacrificed by CO_2_ asphyxiation 15 weeks post diet start. Whole blood was collected from the retro-obital plexus via heparinized capillaries and mixed with EDTA. The mice were then perfused with 10 ml PBS followed by harvest of heart that was immediately embedded into OCT and frozen or fixed in 10% phosphate buffered formalin. Fixed heart was embedded into OCT and sectioned into 10 μm sections. Quantitative assessment of necrotic cores, determined by acellular regions in plaques, and atherosclerotic lesions was performed as described by Baglione and Smith^42^, using the Olympus CX43 Microscope and CellSense software.

### Autophagy assay

RAW-ASC, RAW-ASC-GsdmD^-/-^, RAW Difluo mLC3, HepG2 and THP-1 macrophages were treated ± DSF ranging from 0.5 up to 10 µM, or ± 25 µM Rapamycin (Invivogen; # tlrl-rap) for 2hr. Autophagy markers were probed by microscopy or western blot analysis. Detailed methods are described in *supplementary material and methods* section.

### Western Blotting

Western blot analysis was performed on protein extracts ± various treatment conditions. Detailed methods are described in *supplementary material and methods* section.

### Indirect Immunofluorescence

THP-1 macrophages were treated with indicated dose of DSF and probed for MerTK expression. Detailed methods are described in *supplementary material and methods* section.

### Efferocytosis and phagocytosis assay

THP-1 macrophages were grown in chamber slides and treated ± DSF. Jurkat cells (ATCC) were labeled with 1µM Calcein for 1 hour and washed with PBS 3 times. Calcein labeled cells were treated with 1µM of staurosporine for (16-20) hours to induce apoptosis. After 16 hours of treatment, apoptosis was confirmed by looking at the membrane blebbing of the cells under confocal microscope. Apoptotic Jurkat cells were incubated with THP-1 macrophages in the ratio (1:1) for 4 hours. After 4 hours, cells were washed with 3 times with PBS for 5 minutes each. Then, cells were fixed using 3.7% paraformaldehyde for 30 min. Cells were washed again with PBS and then imaged with confocal microscopy. For phagocytosis assay, the THP-1 macrophages were incubated with Latex beads-rabbit-IgG-FITC complex (Cayman Chemical # 500290) with a final dilution of 1:200 at 37°C for 1 hour. After incubation, cells were washed using an assay buffer and observed under a fluorescence microscope using FITC and DIC channel. Efferocytosis and phagocytosis was determined by measuring mean fluorescence intensity of greater than 30 cells per treatment group.

### Atomic force microscopy (AFM)

THP-1 macrophages treated with ± 5µM DSF for 2h were subjected to AFM analysis as previously described^43^. In brief, a tipless cantilever glued with a 4.5 µm bead was submerged into cell culture where the approach/retraction velocity was 5 µm/s, exerting an indented force of 2nN onto cells. Young’s moduli (modulus of elasticity or stiffness) of the cells were determined using a Hertz model^44^, whereas adhesion, defined as the force required to separate cell surface and cantilever tip, was calculated directly from the force-indentation curves as detailed elsewhere^45,46^.

### Immunofluorescence of aortic sinus

Fresh hearts embedded in OCT were cryosectioned (5 μm) using Leica CM3050 S Cryostat. Sections were fixed in 4% phosphate buffered formalin for 10 minutes at room temperature. After PBS wash, sections were incubated with BLOXALL Blocking Solution (Vector Labs; SP-6000) for 1 hour at room temperature to block non-specific binding. Sections were then incubated overnight with respective primary antibodies at 1:50 dilution After washing, slides were incubated with 1:150 dilution of Alexa-488 goat anti-rabbit secondary (Invitrogen; #A11034) for 1 hour at room temperature. Slide were then washed and mounted with mounting media containing DAPI (Molecular Probes; #36964) and examined using the Nikon Eclipse T*i* confocal and Nikon NIS Elements Imaging Software version 4.13.

### Gut microbiota sequencing

The fresh fecal samples were collected from mice using sterile forceps and then microbial DNA was extracted using the DNeasy PowerSoil ProKit (Qiagen; # 47016), following manufacturer’s instructions. 16S rRNA gene amplicon sequencing and bioinformatics analysis were performed using methods explained earlier^47–49^. Briefly, raw 16S amplicon sequence and metadata, were demultiplexed using split_libraries_fastq.py script implemented in QIIME2^50^. Demultiplexed fastq file was split into sample-specific fastq files using split_sequence_file_on_sample_ids.py script from QIIME2. Individual fastq files without non-biological nucleotides were processed using Divisive Amplicon Denoising Algorithm (DADA) pipeline^51^. The output of the dada2 pipeline (feature table of amplicon sequence variants (an ASV table)) was processed for alpha and beta diversity analysis using phyloseq^52^, and microbiomeSeq (http://www.github.com/umerijaz/microbiomeSeq) packages in R. We analyzed variance (ANOVA) among sample categories while measuring the of α-diversity measures using plot_anova_diversity function in microbiomeSeq package. Permutational multivariate analysis of variance (PERMANOVA) with 999 permutations was performed on all principal coordinates obtained during CCA with the ordination function of the microbiomeSeq package.

### Statistics

Statistics were performed using GraphPad Prism 9. Data is represented as mean ± SD or mean ± SEM for in vitro and in vivo studies, with at least replicates performed for each experiment. For microbiome data, differential abundance analysis was performed using the random-forest algorithm, implemented in the DAtest package (https://github.com/Russel88/DAtest/wiki/usage#typical-workflow). Briefly, differentially abundant methods were compared with False Discovery Rate (FDR), Area Under the (Receiver Operator) Curve (AUC), Empirical power (Power), and False Positive Rate (FPR). Based on the DAtest’s benchmarking, we selected metagenomeSeq and ANOVA as the methods of choice to perform differential abundance analysis. We assessed the statistical significance (p < 0.05) throughout, and whenever necessary, we adjusted P-values for multiple comparisons according to the Benjamini and Hochberg method to control False Discovery Rate^53^. Linear regression (parametric test), and Wilcoxon (Non-parametric) test were performed on genera and ASVs abundances against metadata variables using their base functions in R (version 4.1.2)^54^.

## Results

### Disulfiram inhibits in-vitro and in-vivo Nlrp3 inflammasome activity

GsdmD, the final executor of inflammasome activity, promotes progression of atherosclerosis, with cleaved GsdmD fragment found in human and mice atherosclerotic plaques^14–16^. Disulfiram (DSF) was shown to block GsdmD-mediated pyroptosis and LPS-induced septic death in mice^55^. First, we confirmed that DSF blocks GsdmD activity in macrophages and in mice. WT and GsdmD^-/-^ bone marrow derived macrophages (BMDMs) were primed with LPS then stimulated by ATP. DSF inhibited IL-1β release in WT BMDMs, with no additive effects in GsdmD^-/-^ BMDMs (***Fig. 1a***). For in-vivo efficacy, WT and GsdmD^-/-^ mice (fed with WTD ± 100 mg/kg/day DSF) were induced for inflammasome assembly via LPS+ATP injections as described in our earlier work^14^. DSF-fed mice showed ∼ 80% reduction in IL-1β release in peritoneal lavage (p<0.0001), with no additive effects observed in GsdmD^-/-^ mice (***Fig. 1b***). Dose-response for viability and LDH cytotoxicity assays showed that the 2, 5, and 10 μM of DSF did not affect viability (***Fig. 1c,d***). We chose the 5μM dose for further testing and as shown in ***Fig. 1d***, DSF-treated RAW-ASC cells showed significant reduction (∼88%) in IL-1β release (p<0.0001). As expected, control RAW-ASC-GsdmD^-/-^ cells showed marked reduction in IL-1β release upon LPS+ATP stimulation (***Fig. 1e***) with p<0.0001. These data indicate that Disulfiram inhibits in-vitro and in-vivo Nlrp3 inflammasome activity in a GsdmD-dependent manner.

**Figure 1.**
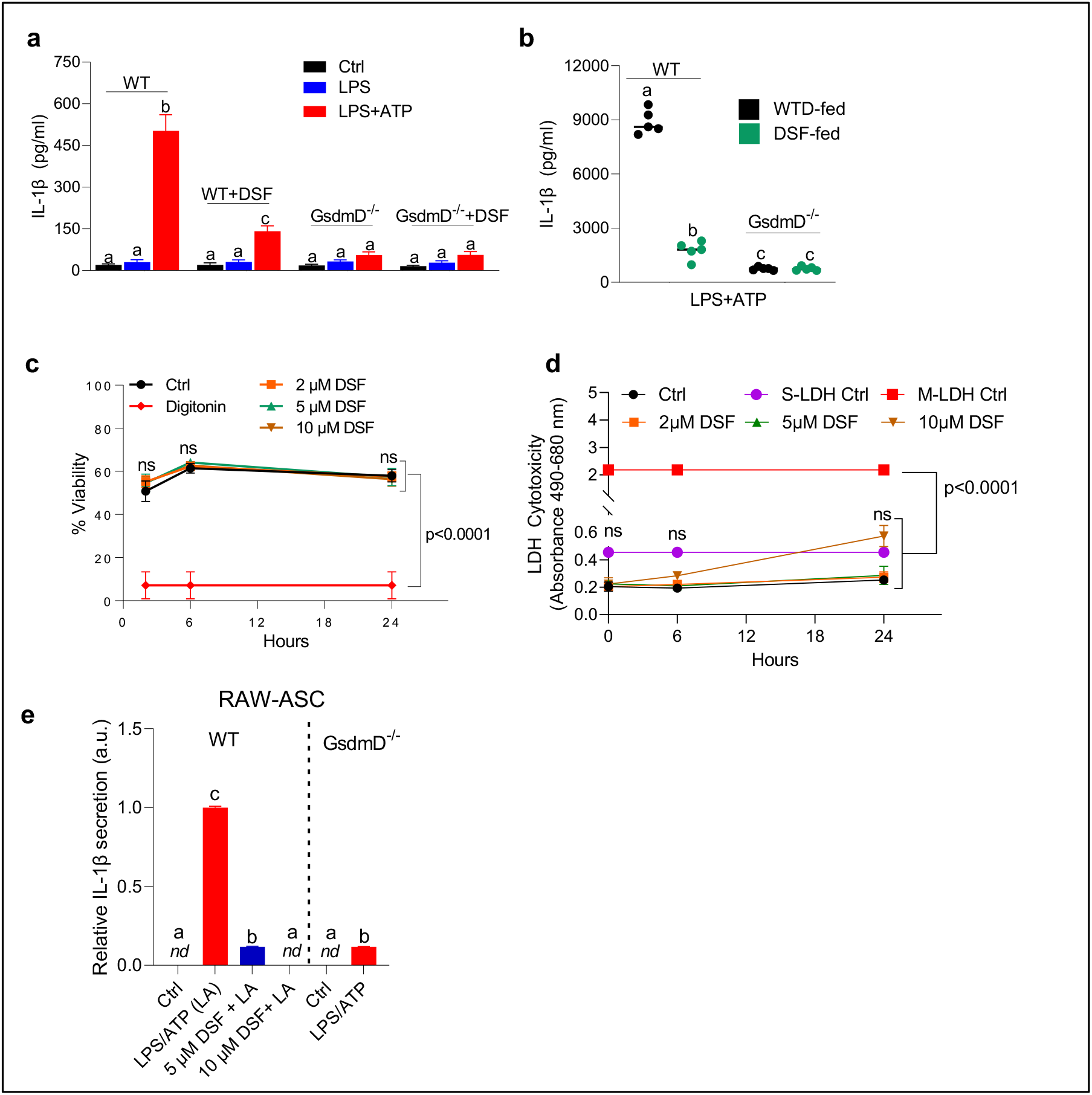
DSF reduces IL-1β secretion in vitro and in-vivo in a GsdmD dependent manner. **a)** ELISA of BMDMs derived from C57BL/6J-WT and C57BL/6J-GsdmD^-/-^ ± DSF treatment ± Nlrp3 inflammasome induction by LPS/ATP; n=5 mice / group. **b)** ELISA of peritoneal lavage from C57BL/6J-WT and C57BL/6J-GsdmD^-/-^ fed with WTD ± DSF diet and induced for inflammasome using LPS+ATP injections; n=5 mice /group. **c)** Live/dead fixable blue dead cell assay for cell viability of RAW-ASC cells treated with 2, 5, and 10 μM DSF for 2, 6, and 24h.; n= 4 **d)** CyQuant LDH cytotoxicity assay for toxicity of RAW-ASC cells treated with 2, 5, and 10 μM DSF for 0, 6, and 24h. Spontaneous LDH release (S-LDH) and positive control Maximum LDH release (M-LDH); n=3. **e)** ELISA for IL-1β from WT RAW-ASC and RAW-ASC-GsdmD^-/-^ cells treated with ± 5 and 10μM DSF and primed with 1 μg/ml LPS for 4h, followed by stimulation with 1mM ATP for 30 minutes. Data is plotted relative to WT RAW-ASC cells treated with LPS + ATP; n=3. Significance was determined by one-way ANOVA with different letters indicating significant difference of p<0.05, groups sharing same letters indicate no significant difference; ns= non significant, nd = not detectable.

### Disulfiram reduces atherosclerosis in hyperlipidemic mice

Previous studies demonstrated that global knockout of GsdmD resulted in smaller atherosclerotic lesions in hyperlipidemic mice^14,16^. To determine if DSF can reduce atherosclerosis, apoE^-/-^ mice were fed an atherogenic western type diet (WTD) or WTD containing 100mg/kg/day DSF for 15 weeks, as shown in schematic diagram (***Fig. S1a***). DSF-fed male mice showed a small but significant increase in food intake vs. WTD-fed males (p=0.035), while no differences were found in female mice (p=0.909) (***Fig. S1b***). There were no significant changes in body weight in DSF-fed mice in both sexes (p=0.154 for males, p=0.094 for females, ***Fig. S1c***), nor were there any significant changes in liver weight in DSF-fed vs. WTD-fed control mice (p=0.681 for males, 0.269 for females, ***Fig. S1d***). The DSF-fed mice showed significantly increased plasma cholesterol (p=0.012 for males, and p=0.009 for females (***Fig. S1e***). Plasma was fractionated using FPLC and the peaks showed no differences in distribution of apolipoprotein B particles (VLDL, LDL, IDL) or HDL-cholesterol (p=0.731 for males, and p=0.682 for females) (***Fig. S1f***). We also determined if DSF altered complete blood counts in these hyperlipidemic mice, but no significant differences were found in number of platelets, white blood cells, eosinophils, monocytes, lymphocytes, or neutrophils (***Fig. S2*)**. Red blood cells showed no difference in males fed with DSF vs. WTD only, but there was a small but significant decrease in number of RBC in females (p=0.004, ***Fig. S2***).

The progression of atherosclerosis was determined by the oil red O staining of aortic root sections from WTD-fed vs. DSF-fed mice. As shown in ***Fig. 2a,b***, atherosclerotic lesions were significantly smaller with 26.7% and 29.4% reduction in males and females, respectively (p=0.008 for males, and p=0.005 for females). As shown in ***Fig. 2a,c***, necrotic cores were also significantly decreased in DSF vs. WTD-fed mice, with 49.85% and 45.63% reduction in males and females, respectively (p=0.0006 for males and p=0.0208 for females). These data indicate that DSF reduces atherosclerosis independent of cholesterol levels and without altering host immune cell profile.

**Figure 2.**
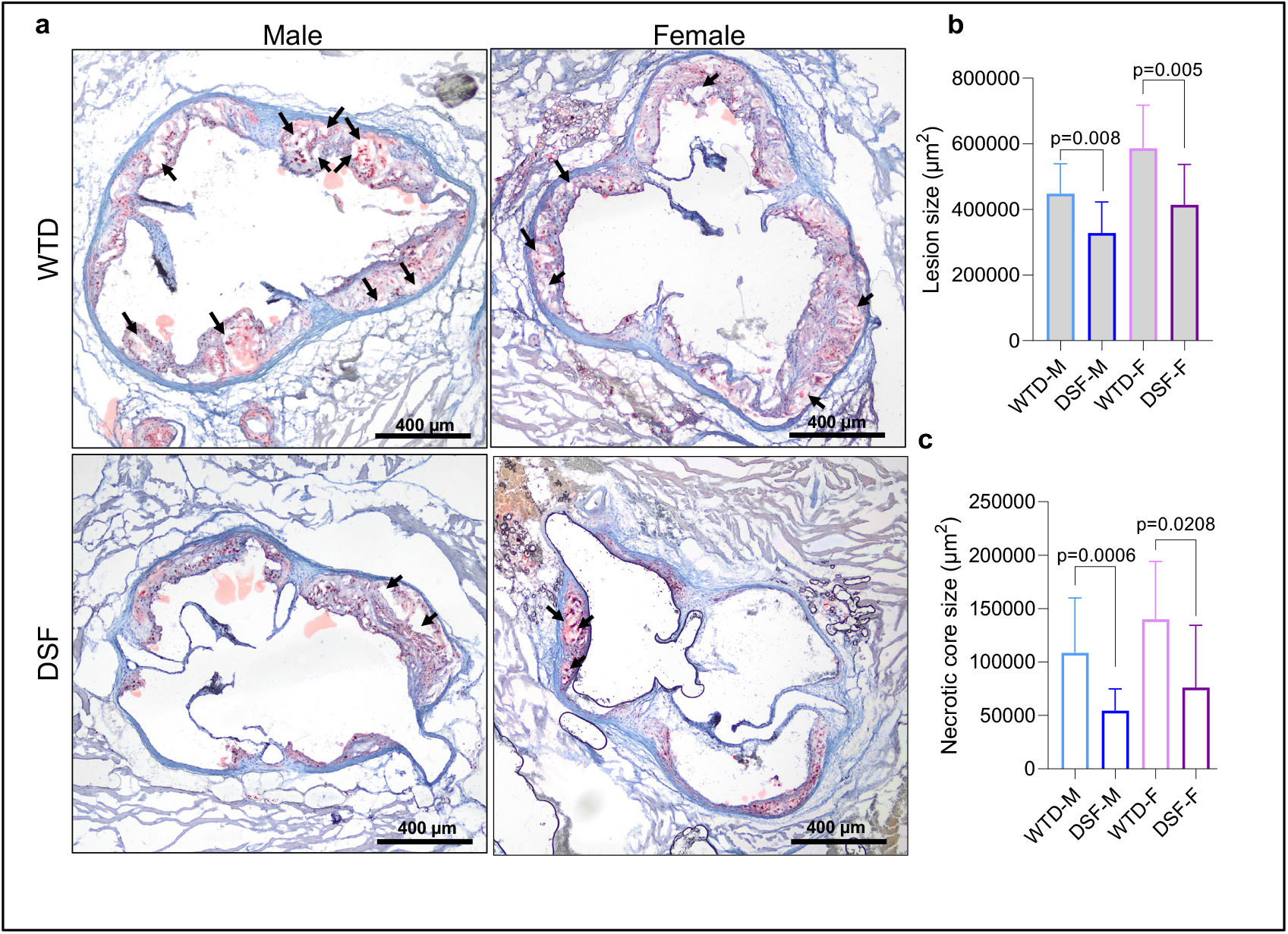
DSF reduces atherosclerotic lesions and necrotic cores in hyperlipidemic apoE^-/-^ mice. **a)** Oil red O staining of atherosclerotic lesions in aortic sinus root sections (10μm) from mice fed with WTD ± DSF; n=9-13 mice per group and per sex, p-values determined by t-test. **b)** Quantification of atherosclerotic lesions. **c)** Quantification of necrotic core in atherosclerotic lesions (mean ± SD; p-values determined by t-test; n=9-13 mice per group and per sex). Black arrows show necrotic core area.

### Disulfiram induces autophagy in-vitro and in-vivo

Autophagy plays an atheroprotective role, but is defective in advanced atherosclerotic lesions, and can serve as target for anti-atherosclerotic therapeutic^3,25,26,28^. To determine if DSF can induce autophagy, we treated RAW-Difluo mLC3 reporter cells (Invivogen) with DSF or rapamycin (positive control). RAW-Difluo mLC3 cells express the RFP::GFP::LC3 fusion protein where RFP (acid-stable) and GFP (acid-sensitive) and colocalization of GFP and RFP is signal for increased autophagosome formation. DSF treatment increased the number and colocalization of LC3-GFP and LC3-RFP puncta vs. control cells (***Fig. 3a,b***), indicating autophagosome formation. This data indicates fusion of autophagosomes with lysosomes and increased autophagy flux in DSF treated cells. We confirmed DSF-induced autophagy in RAW-ASC cells, by probing for autophagy markers p62 and LC3-II. The adaptor protein p62 binds to unfolded proteins and damaged membranes to chaperone them to the autophagophore for degradation, thus p62 levels are also reduced during autophagy induction. As shown in ***Fig. 3c,d*** and ***Fig. S3***, DSF-treated cells showed reduced p62 (p=0.005) and increased LC3-II (p=0.002). These results were also confirmed in THP-1 macrophages, where LC3-II levels were increased upon DSF-treatment (***Fig. S4***). As DSF is a potent inhibitor of GsdmD, we determined if DSF effects on autophagy were via GsdmD blockage. Interestingly, the autophagy induction by DSF was independent of GsdmD, as autophagy was induced in GsdmD^-/-^ cells treated with DSF (***Fig. 3e, Fig. S3***). Next, we determined if DSF could induce autophagy in-vivo. The heart sections containing atherosclerotic lesions from mice fed with WTD ± DSF were probed with anti-mouse specific anti-LC3 antibody. As shown in ***Fig. 4a,b***, expression of LC3 was significantly higher in atherosclerotic lesion from DSF-fed vs. WTD-fed control mice (p<0.0001). To determine if DSF induced autophagy in other tissues, liver homogenates from mice fed with WTD ± DSF were probed for LC3-II expression. As shown in ***Fig. 4c,d*** and ***Fig. S5***, expression of LC3-II was significantly higher in liver of DSF-fed vs. WTD-fed control mice (p=0.015 for males and p=0.041 for females). Autophagy is also known to reduce lipid-droplets in liver^56,57^. Thus, to determine if autophagy induction in liver alters accumulation of lipid-droplets, the liver sections from mice fed with WTD ± DSF were stained with H & E stain. The WTD-fed mice showed extensive presence of lipid-droplets in liver, while DSF-fed mice had significantly lower accumulation of lipid droplets (***Fig. 4e***). To further confirm effect of DSF on autophagy in liver cells, we treated human hepatocytes (HepG2) with various doses of DSF and probed for LC3-II expression. As shown in ***Fig. 4f,g*** and ***Fig. S5***, DSF induced LC3-II expression in a dose-dependent manner (p=0.01 for 5μM and 10μM).

**Figure 3.**
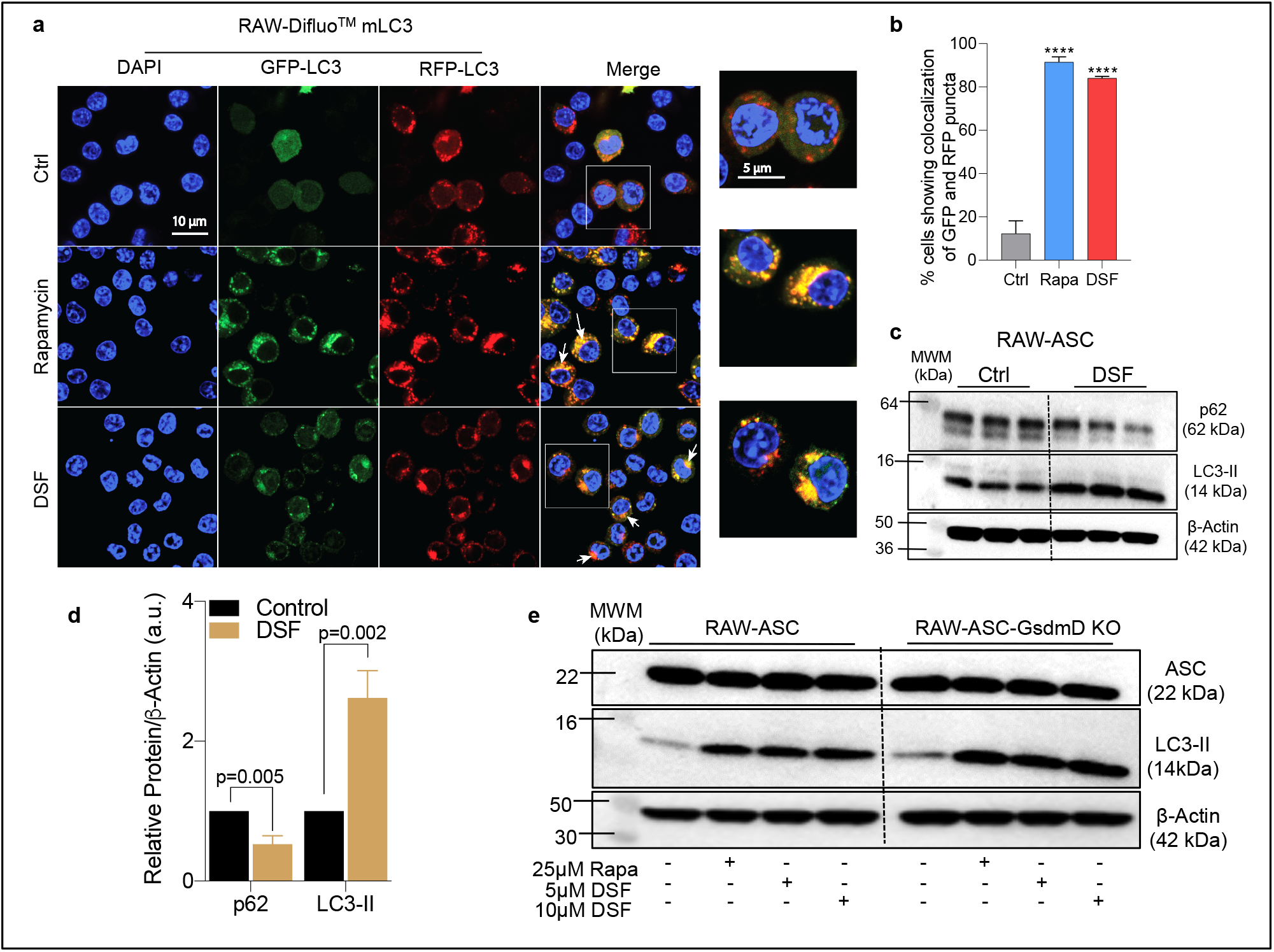
DSF induces autophagy in macrophages independent of GsdmD. RAW-Difluo mLC3 (autophagy reporter), RAW-ASC, and RAW-ASC-GsdmD^-/-^, cells were treated ± 5μM DSF for 2h. **a)** Microscopy of autophagy reporter cells line RAW-Difluo mLC3 ± DSF and ± positive control rapamycin (Rapa). **b)** % cells showing colocalization of GFP-LC3 and RFP-LC3 puncta in RAW-Difluo mLC3 ± DSF treatment n=100 cells/group; ****p=0.0001 determined by t-test; **c)** Western blot and **d)** quantification of autophagy markers p62 and LC3-II in RAW-ASC cells, n=3, p-values determined by t-test. **e**) Western blot of autophagy markers LC3-II in RAW-ASC vs. RAW-ASC-GsdmD^-/-^ cells ± 5 μM or 10 μM DSF ± positive control rapamycin (Rapa).

**Figure 4.**
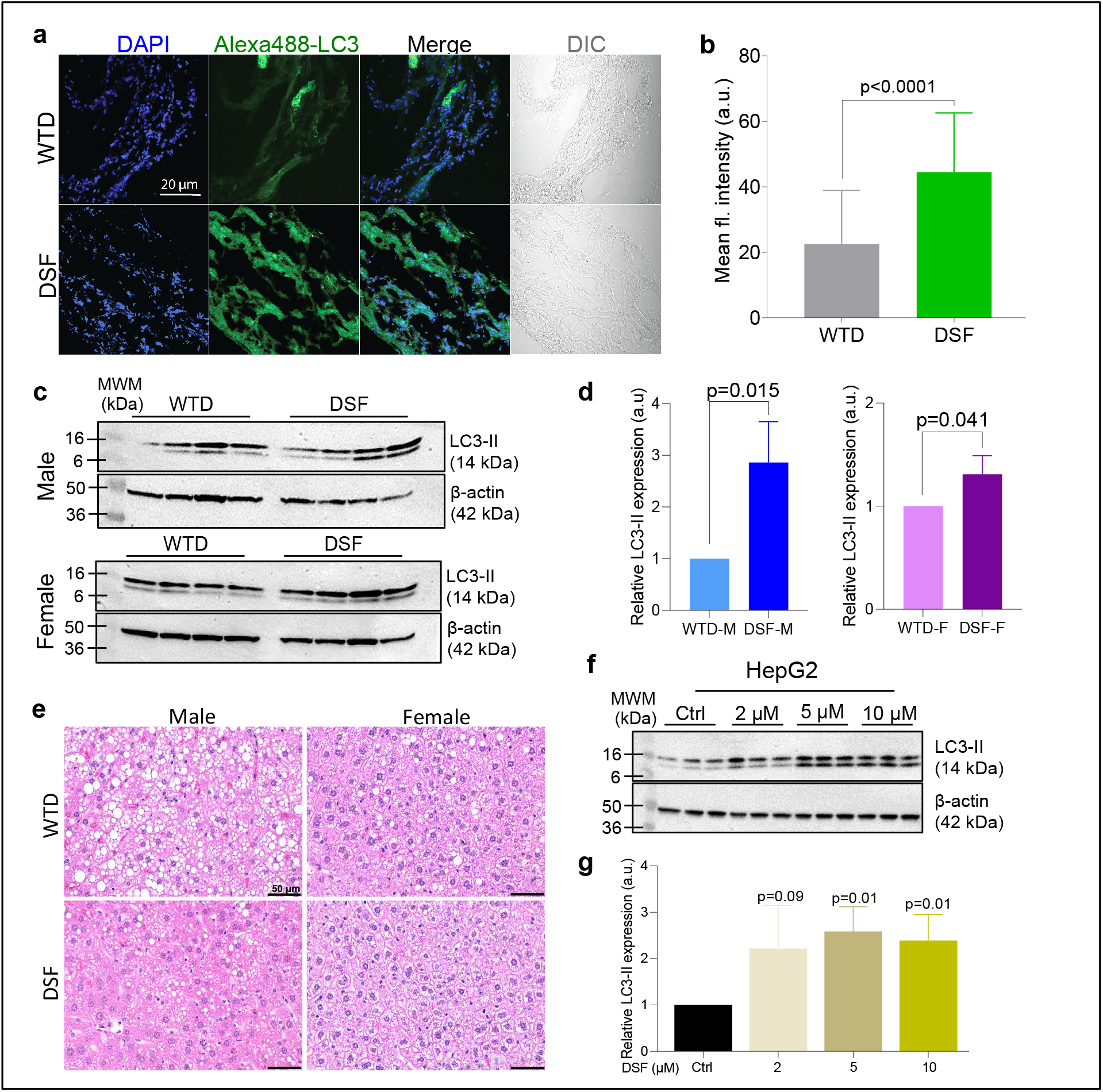
DSF induces autophagy in atherosclerotic plaques/liver tissue of hyperlipidemic apoE^-/-^ mice, and in cultured hepatocytes. **a)** Expression of LC3 in atherosclerotic lesions by indirect immunofluorescence microscopy using LC3 antibody and Alexa-488 labeled secondary antibody. **b)** Quantification of mean fluorescence intensity, n=50 fields; p-value determined by t-test. **c)** Western blot of LC3-II in liver homogenate of mice fed with WTD ± DSF, upper panel shows males and lower panel shows females; n=4. **d)** Quantification of LC3-II western blot band intensity in male and female mice, n=4, p-values determined by t-test. **e)** Representative H & E staining of liver sections from hyperlipidemic mice fed with WTD ± DSF for 15 weeks. **f)** Western blot of autophagy marker LC3-II in HepG2 cells treated with various doses of DSF vs. untreated cells, n=3. **g)** Quantification of LC3-II western blot band intensity ± SD); n=3, p-values determined by t-test.

### Disulfiram increases efferocytosis and phagocytosis in macrophages

Effect of DSF on the atheroprotective pathways such as phagocytosis^58^ and efferocytosis^4,32^ was determined. Phagocytosis in DSF-treated macrophages was probed by incubating cells with IgG-FITC coated latex beads. As shown in ***Fig. 5a,b*** *and **Fig. S6**,* DSF-treated THP-1 macrophages showed a robust increase in phagocytic activity vs. control (p=0.005), as determined by uptake of FITC-coated latex beads. As shown in ***Fig. 5c,d*** *and **Fig. S6**,* DSF-treated THP-1 macrophages also showed markedly enhanced efferocytosis vs. control (p=0.015), as determined by the uptake of calcein-labeled apoptotic Jurkat cells by macrophages. To decipher the mechanism of enhanced efferocytosis, effect of DSF on expression of efferocytic receptor MerTK was determined. DSF-treated THP-1 macrophages showed markedly enhanced cell-surface expression of MerTK (***Fig. 5e***). As phagocytosis and efferocytosis are effected by biophysical properties of macrophage cell membrane^43,59,60^, we measured elasticity and adhesion of DSF-treated vs. control cell by Atomic force microscopy (AFM). As shown in ***Fig. 5f***, DSF treatment increased membrane rigidity of THP-1 macrophages, denoted by a significant increase in Young’s Modulus of elasticity (p=0.0007). Additionally, AFM showed an increase in cell adhesion force in DSF-treated macrophages vs. non-treated controls (***Fig. 5g***, p<0.0001).

**Figure 5.**
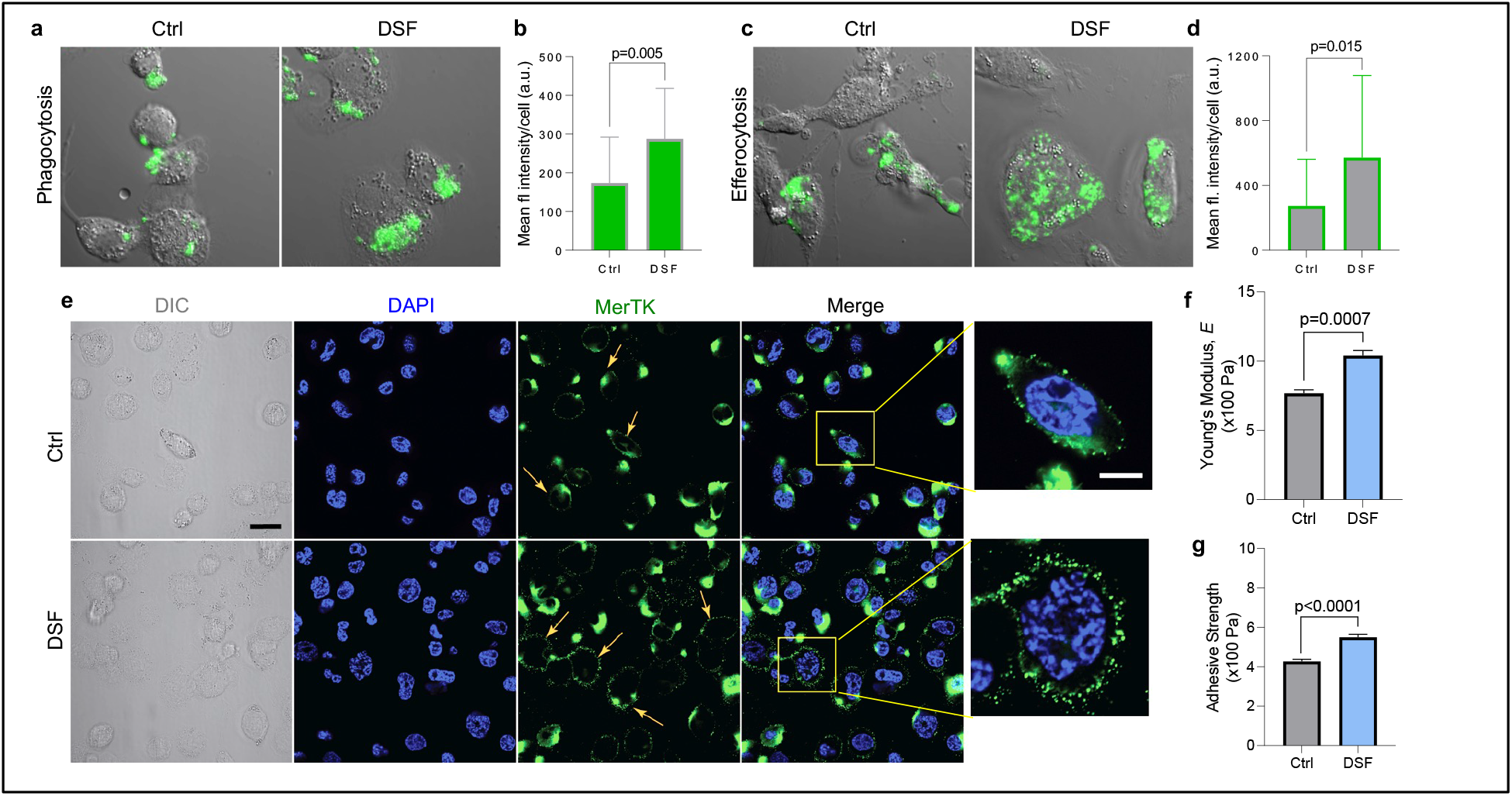
DSF enhances efferocytic and phagocytic activity of macrophages. THP −1 macrophages pretreated with ± 5µM DSF for 2h, were incubated with either IgG-FITC coated-latex beads for 1hr (phagocytosis study) or calcein-labeled staurosporine-induced apoptotic Jurkat cells for 4hr (efferocytosis study). **a)** Microscopy and **b)** quantification of phagocytosis of FITC-labeled latex beads, with n ≥ 20 cells per treatment group, with p-values determined by t-test. **c)** Microscopy and **d)** quantification of efferocytosis of calcein-labeled apoptotic Jurkat cells, with n ≥ 20 cells per treatment group, with p-values determined by t-test. **e)** Indirect immunofluorescence of MerTK in THP-1 macrophages ± 5µM DSF for 2h, using anti-human MerTK primary antibody and Alexa488-labeled goat anti-rabbit secondary antibody, with cells imaged by confocal microscopy; black scale bar is 25 µm and white scale bar is 10µm. **f)** Young’s modulus of elasticity by AFM and **g)** Adhesion strength by AFM; 30 cells analyzed per group with p-values determine by t-test.

### DSF treated hyperlipidemic mice showed altered gut microbiota profile

The gut microbiome plays a pivotal role in the progression of atherosclerosis^5,34,61,62^. DSF has been shown to have antimicrobial effects^24,63^. Thus, we determined if DSF could modulate gut microbiome of hyperlipidemic apoE^-/-^ mice. The 16s ribosomal RNA (16s rRNA)-based qPCR sequencing was performed on the fresh fecal samples from apoE^-/-^ mice fed with WTD ± DSF diet for 15 weeks. The alpha and beta diversity in microbial population across each group was analyzed. Alpha diversity is a measure of microbiome diversity/complexity in each sample, whereas the beta diversity is a measure of similarity or dissimilarity between groups. As shown in ***Fig. 6***, the gut microbiome profile was significantly different between the DSF-fed vs. WTD-fed control mice, with alterations seen in both alpha (p<0.05) and beta (p<0.05) diversity (***Fig. 6a,b***). The DSF-fed mice showed marked reduction in levels of *Romboutsia, Lactococcus,* and *Blautia* bacteria genera compared to that of WTD-fed mice (p<0.05) and highly significant enrichment in *Turicibacter, Bidifobacterium, Lactobacillus, and Akkermansia,*(p<0.05 for all species, ***Fig. 6c,d***). Collectively, these data indicate that DSF modulates multiple atheroprotective pathways and its effect on atherosclerosis may be mediated via GsdmD-dependent as well as in GsdmD-independent manner.

**Figure 6.**
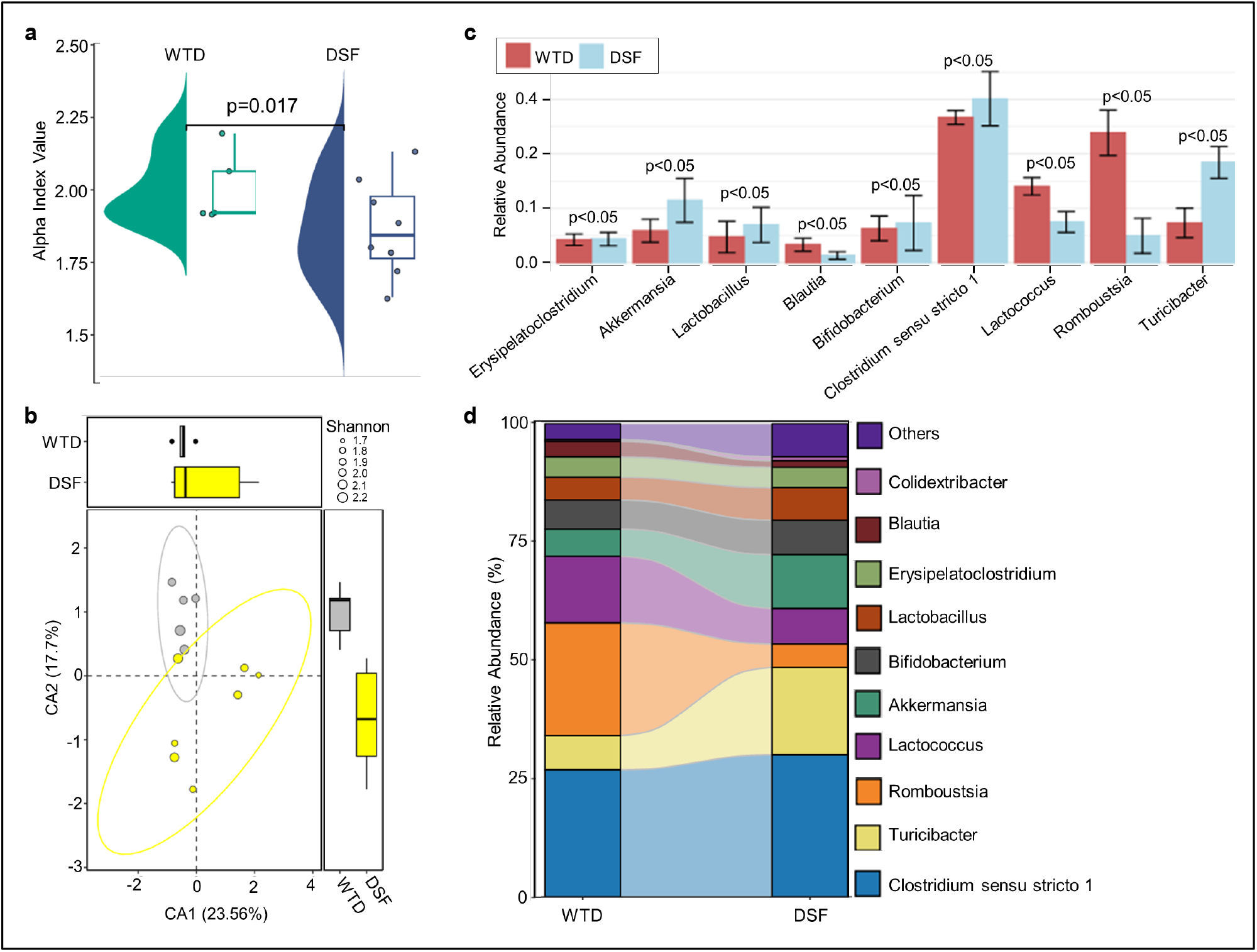
DSF modulates gut microbiota in hyperlipidemic apoE^-/-^ mice. The gut microbiota profile of apoE^-/-^ mice fed with WTD ± DSF for 15 weeks using 16sRNA-based qPCR sequencing; n=5 for WTD and n=8 for DSF **a, b)** Shannon plots showing alpha diversity of gut microbiota in WTD vs. DSF fed mice. **c)** Graph showing relative abundance of bacterial genera that are significantly altered in WTD vs. DSF fed mice; p<0.05. **d)** Relative abundance of various bacterial species in WTD vs. DSF-fed mice.

## Discussion

Low-density lipoprotein-cholesterol (LDL-C) lowering therapeutics provide irrefutable benefits for treating CVD, but there is still a 50 to 70% residual risk for the major adverse coronary event (MACE) even in high-dose statin-treated subjects. Thus, adjuvant therapies targeting pathways other than lowering LDL-C are needed. The Canakinumab Anti-Inflammatory Thrombosis Outcome Study (CANTOS) trial, using interleukin-1 beta (IL-1β) antibody in humans, showed the feasibility of adjuvant therapies that target inflammation, independent of lipid levels. But in 2018, the FDA declined to approve Canakinumab (Ilaris) for cardiovascular risk reduction on the strength of data. Thus, a drug that targets multiple pathways simultaneously may serve as a adjuvant therapeutic for chronic and complex disease such as atherosclerosis. Blocking pyroptosis, rather than letting cells rupture and targeting only released IL-1β, may be more beneficial as other inflammatory molecules such as IL-18 and High-mobility group box protein 1 (Hmgb1) can also be blocked. Moreover, earlier work from our lab and others showed that blocking pyroptosis shifts the balance toward apoptotic cell death^14,16^. Furthermore, a previous study showed that DSF can also induce ER stress and unfolded protein response-mediated apoptosis^64^. Apoptosis is beneficial in dampening atherosclerosis and cells undergoing apoptosis can be efficiently cleared via phagocytosis and efferocytosis^29^. In addition, inducing autophagy in plaques may result in degradation of lipid droplets and generation of free cholesterol that can be effluxed from arterial foam cells via apoA1-ABCA1 pathway^65–67^. In addition to targeting cellular pathways, modulation of gut microbiota to promote enrichment of atheroprotective and depletion of atherogenic bacterial species could be another strategy to slow down atherosclerosis progression^34,68^.

Given the recent findings showing the role of GsdmD in atherosclerosis^13–16^, we tested the anti-atherosclerotic activity and effect of GsdmD inhibitor, Disulfiram (DSF), on atherosclerosis-related pathways. DSF, also known by its trade name Antabuse, has been approved by the FDA and has been in use for decades for treating chronic alcoholism, as an inhibitor of alcohol metabolizing enzyme aldehyde dehydrogenase (ALDH). DSF effects seem to be broader than targeting ALDH alone, as highlighted by several clinical trials that are underway for repurposing DSF for various diseases (https://clinicaltrials.gov/search?intr=Disulfiram), but it’s efficacy as an anti-atherosclerosis agent is not yet explored. We found that DSF reduced atherosclerosis progression in a hyperlipidemic apoE^-/-^ mouse model and modulated various pathways involved in atherosclerosis.

Activation of the Nlrp3 inflammasome in advanced human atherosclerotic plaques results in GsdmD cleavage and further disease progression with thinning of the fibrous cap and increase in the necrotic core area in plaques^10,12^. In agreement with a previous study^55^, DSF-treated macrophages and DSF-fed hyperlipidemic mice showed reduced IL-1β release upon Nlrp3 inflammasome assembly (***Fig. 1***), indicating reduction in pore formation. DSF does not block Nlrp3 inflammasome directly, nor does it blocks cleavage of GsdmD, but rather it inhibits the terminal event of membrane pore formation and IL-1β release^55^. Thus DSF-treated cells are expected to retain their ability to destroy intracellular pathogens. DSF, thus may serve as better therapy for atherosclerosis than Nlrp3 inhibitors, as Nlrp3 is essential for countering several pathogens^69^. Using a hyperlipidemic apoE^-/-^ mouse model of atherosclerosis, we found that DSF reduced size of atherosclerotic lesions (***Fig. 2***). The effect of atherosclerosis is not attributable to feeding behavior, as food-intake was not compromised as the DSF-fed male mice ate slightly more food vs. WTD-fed control (***Fig. S1***). The DSF effects on atherosclerosis were independent of plasma cholesterol levels as there was a small increase in plasma cholesterol in DSF-fed mice (***Fig. S1***). DSF-fed mice did not show any significant changes in plasma levels of platelets, white blood cells, eosinophils, monocytes, lymphocytes, and neutrophils (***Fig. S2***). These data indicate that DSF effects on atherosclerosis are unlikely due to major changes the in blood cholesterol or immune cell population. It is more probable that the effects are due to changes in the intra-plaque immune cell profile, with expected lower inflammation and blunted recruitment of monocytes in plaque area in DSF vs. WTD-fed control mice. In addition to apoptotic cell death, the vast majority of cells in advanced lesions undergo necrotic cell death^70^, leading to further amplification in inflammation. The role of the necrotic core in pathogenesis of atherosclerosis is not fully deciphered, but it is widely believed to stimulate inflammatory pathways and reduce plaque stability^70^. DSF-fed mice showed reduced necrotic core within atherosclerotic plaques (***Fig. 2***), indicating that DSF can also modulate necrosis in plaques. DSF may be reducing necrosis via reducing intra-plaque IL-1β levels, or by enhancing phagocytic and efferocytic activity of macrophages. Both phagocytosis and efferocytosis are essential for blocking progression of atherosclerosis^30,58^. Efferocytosis is regulated by interaction between exposed phosphatidylserine (PS) on apoptotic cells and PS receptors (MerTK or AxlTK) on macrophages^4,32^. DSF enhanced expression of MerTK (***Fig. 5***), which may result in efficient clearance of necrotic cells from atherosclerotic plaques. DSF also up regulated autophagy, a major atheroprotective pathway. DSF induced autophagy in macrophages (***Fig. 3***) and DSF-fed mice also showed induced autophagy in atherosclerotic plaques DSF-fed mice showed increased expression of LC3 in plaques (***Fig. 4***), indicative of induced autophagosomes formation. The liver homogenate of DSF-fed mice also showed increased LC3-II expression vs. WTD-fed control mice. Similar results were found in hepatocytes (***Fig. 4***), indicating that DSF effects on autophagy induction may be global, and not limited to arterial macrophages. The liver sections from DSF-fed mice also showed fewer lipid droplets vs. WTD-fed control mice (***Fig. 4***), and one of the mechanisms for this effect could be DSF-induced lipophagy in liver. However, the effect of increased lipid degradation in liver did not get translated into lower plasma cholesterol levels and we believe that this is due to inability of apoE^-/-^ mice to clear cholesterol from systemic circulation. In addition to testing DSF effects on cellular atheroprotective pathways, we also profiled gut microbiome of DSF-fed mice. The rationale was that recent studies showed cross talk between GsdmD and gut microbiota^71,72^ and GsdmD is functionally deactivated in DSF-fed mice. The DSF-fed hyperlipidemic mice showed markedly altered gut microbiota profile, with reduction in levels of *Romboutsia, Lactococcus,* and *Blautia* bacteria while bacteria species such as *Turicibacter, Bidifobacterium, Lactobacillus, and Akkermansia* were enriched compared to WTD-fed controls (***Fig. 6***). Previous studies have shown atherogenic role of *Romboutsia*^62,73^, and atheroprotective role of *Akkermansia*^62,68,73^ *Bifidobacterium*^73,74^, *Lactobacillus*^74^ and *Turicibacter*^75^ in hyperlipidemic apoE^-/-^ mouse model of atherosclerosis.

Based on our data, DSF can serve as an adjuvant therapy for treating atherosclerosis and CVD. The DSF effects on atherosclerosis seem to be GsdmD-dependent (IL-1β release) as well as GsdmD-independent (autophagy). Thus, further work is required to understand the pleiotropic effects of DSF on atherosclerosis related pathways and gut microbiota. For example, microbial transplantation using germ-free mice and bone marrow transplantations are needed to decipher the contribution of gut microbiota and myeloid cells on DSF mediated modulation of atherosclerosis. Furthermore, knockouts of Atg7 or MerTK in GsdmD^-/-^ background can provide evidence for role of DSF-induced autophagy and efferocytosis in atherosclerosis. An elegant study showed DSF can also prevent body weight gain in mice fed with obesogenic diet and reversed diet-induced obesity^41^. Obesity is well known risk factor for accelerated atherosclerosis and increased rates of cardiovascular death^76,77^. Thus, DSF can be repurposed as an adjuvant therapy for chronic inflammatory metabolic diseases such as atherosclerosis and obesity. Further work is required to delineate the specific contributions of various pathways toward anti-CVD properties of DSF.

## Acknowledgements

This work is supported by NIH R01-HL148158, American Heart Association Transformational project (AHA-TP) award 23TPA1063910, and Cleveland State University start-up funds to K.G. We acknowledge help in data acquisition and data analysis from Dr. Naseer Sangwan from Cleveland Clinic Microbiome Core for 16s rRNA sequencing of gut microbiota.

## Author Contributions

KG designed and directed the research. KG and CAT designed experiments. CAT, KT AP, NB, MRK, EO, KN, EO, GB, EGE and KG performed experiments and analyzed the data. JDS, KN, PN, and CRK provided material support and data analysis. CAT and KG drafted the manuscript. All authors critically reviewed the manuscript.

## References

1. Libby P. Inflammation in Atherosclerosis-No Longer a Theory. Clin Chem. 2021;67:131–142. doi: 10.1093/clinchem/hvaa275

2. Westerterp M, Fotakis P, Ouimet M, Bochem AE, Zhang H, Molusky MM, Wang W, Abramowicz S, la Bastide-van Gemert S, Wang N, et al. Cholesterol Efflux Pathways Suppress Inflammasome Activation, NETosis, and Atherogenesis. Circulation. 2018;138:898–912. doi: 10.1161/CIRCULATIONAHA.117.032636

3. Ouimet M, Ediriweera H, Afonso MS, Ramkhelawon B, Singaravelu R, Liao X, Bandler RC, Rahman K, Fisher EA, Rayner KJ, et al. microRNA-33 Regulates Macrophage Autophagy in Atherosclerosis. Arterioscler Thromb Vasc Biol. 2017;37:1058–1067. doi: 10.1161/ATVBAHA.116.308916

4. Doran AC, Yurdagul A, Jr., Tabas I. Efferocytosis in health and disease. Nat Rev Immunol. 2020;20:254-267. doi: 10.1038/s41577-019-0240-6

5. Brown JM, Hazen SL. Microbial modulation of cardiovascular disease. Nat Rev Microbiol. 2018;16:171–181. doi: 10.1038/nrmicro.2017.149

6. Hassan M. CANTOS: A breakthrough that proves the inflammatory hypothesis of atherosclerosis. Glob Cardiol Sci Pract. 2018;2018:2. doi: 10.21542/gcsp.2018.2

7. Libby P. Interleukin-1 Beta as a Target for Atherosclerosis Therapy: Biological Basis of CANTOS and Beyond. J Am Coll Cardiol. 2017;70:2278–2289. doi: 10.1016/j.jacc.2017.09.028

8. Ridker PM, Libby P, MacFadyen JG, Thuren T, Ballantyne C, Fonseca F, Koenig W, Shimokawa H, Everett BM, Glynn RJ. Modulation of the interleukin-6 signalling pathway and incidence rates of atherosclerotic events and all-cause mortality: analyses from the Canakinumab Anti-Inflammatory Thrombosis Outcomes Study (CANTOS). Eur Heart J. 2018;39:3499–3507. doi: 10.1093/eurheartj/ehy310

9. Soehnlein O, Libby P. Targeting inflammation in atherosclerosis -from experimental insights to the clinic. Nat Rev Drug Discov. 2021;20:589–610. doi: 10.1038/s41573-021-00198-1

10. Duewell P, Kono H, Rayner KJ, Sirois CM, Vladimer G, Bauernfeind FG, Abela GS, Franchi L, Nunez G, Schnurr M, et al. NLRP3 inflammasomes are required for atherogenesis and activated by cholesterol crystals. Nature. 2010;464:1357–1361. doi: 10.1038/nature08938

11. Chu J, Thomas LM, Watkins SC, Franchi L, Nunez G, Salter RD. Cholesterol-dependent cytolysins induce rapid release of mature IL-1beta from murine macrophages in a NLRP3 inflammasome and cathepsin B-dependent manner. J Leukoc Biol. 2009;86:1227–1238. doi: 10.1189/jlb.0309164

12. Rajamaki K, Lappalainen J, Oorni K, Valimaki E, Matikainen S, Kovanen PT, Eklund KK. Cholesterol crystals activate the NLRP3 inflammasome in human macrophages: a novel link between cholesterol metabolism and inflammation. PLoS One. 2010;5:e11765. doi: 10.1371/journal.pone.0011765

13. Fidler TP, Xue C, Yalcinkaya M, Hardaway B, Abramowicz S, Xiao T, Liu W, Thomas DG, Hajebrahimi MA, Pircher J, et al. The AIM2 inflammasome exacerbates atherosclerosis in clonal haematopoiesis. Nature. 2021;592:296–301. doi: 10.1038/s41586-021-03341-5

14. Opoku E, Traughber CA, Zhang D, Iacano AJ, Khan M, Han J, Smith JD, Gulshan K. Gasdermin D Mediates Inflammation-Induced Defects in Reverse Cholesterol Transport and Promotes Atherosclerosis. Front Cell Dev Biol. 2021;9:715211. doi: 10.3389/fcell.2021.715211

15. Jiang M, Sun X, Liu S, Tang Y, Shi Y, Bai Y, Wang Y, Yang Q, Yang Q, Jiang W, et al. Caspase-11-Gasdermin D-Mediated Pyroptosis Is Involved in the Pathogenesis of Atherosclerosis. Front Pharmacol. 2021;12:657486. doi: 10.3389/fphar.2021.657486

16. Puylaert P, Van Praet M, Vaes F, Neutel CHG, Roth L, Guns PJ, De Meyer GRY, Martinet W. Gasdermin D Deficiency Limits the Transition of Atherosclerotic Plaques to an Inflammatory Phenotype in ApoE Knock-Out Mice. Biomedicines. 2022;10. doi: 10.3390/biomedicines10051171

17. Shi H, Gao Y, Dong Z, Yang J, Gao R, Li X, Zhang S, Ma L, Sun X, Wang Z, et al. GSDMD-Mediated Cardiomyocyte Pyroptosis Promotes Myocardial I/R Injury. Circ Res. 2021;129:383–396. doi: 10.1161/CIRCRESAHA.120.318629

18. de Sa KSG, Amaral LA, Rodrigues TS, Ishimoto AY, de Andrade WAC, de Almeida L, Freitas-Castro F, Batah SS, Oliveira SC, Pastorello MT, et al. Gasdermin-D activation promotes NLRP3 activation and host resistance to Leishmania infection. Nat Commun. 2023;14:1049. doi: 10.1038/s41467-023-36626-6

19. He WT, Wan H, Hu L, Chen P, Wang X, Huang Z, Yang ZH, Zhong CQ, Han J. Gasdermin D is an executor of pyroptosis and required for interleukin-1beta secretion. Cell Res. 2015;25:1285–1298. doi: 10.1038/cr.2015.139

20. Man SM, Kanneganti TD. Gasdermin D: the long-awaited executioner of pyroptosis. Cell Res. 2015;25:1183–1184. doi: 10.1038/cr.2015.124

21. Kayagaki N, Stowe IB, Lee BL, O’Rourke K, Anderson K, Warming S, Cuellar T, Haley B, Roose-Girma M, Phung QT, et al. Caspase-11 cleaves gasdermin D for non-canonical inflammasome signalling. Nature. 2015;526:666–671. doi: 10.1038/nature15541

22. Ding J, Wang K, Liu W, She Y, Sun Q, Shi J, Sun H, Wang DC, Shao F. Pore-forming activity and structural autoinhibition of the gasdermin family. Nature. 2016;535:111–116. doi: 10.1038/nature18590

23. Mozzini C, Pagani M. Cardiovascular Diseases: Consider Netosis. Curr Probl Cardiol. 2021:100929. doi: 10.1016/j.cpcardiol.2021.100929

24. Adrover JM, Carrau L, Dassler-Plenker J, Bram Y, Chandar V, Houghton S, Redmond D, Merrill JR, Shevik M, tenOever BR, et al. Disulfiram inhibits neutrophil extracellular trap formation and protects rodents from acute lung injury and SARS-CoV-2 infection. JCI Insight. 2022;7. doi: 10.1172/jci.insight.157342

25. Liao X, Sluimer JC, Wang Y, Subramanian M, Brown K, Pattison JS, Robbins J, Martinez J, Tabas I. Macrophage autophagy plays a protective role in advanced atherosclerosis. Cell Metab. 2012;15:545–553. doi: 10.1016/j.cmet.2012.01.022

26. Sergin I, Evans TD, Zhang X, Bhattacharya S, Stokes CJ, Song E, Ali S, Dehestani B, Holloway KB, Micevych PS, et al. Exploiting macrophage autophagy-lysosomal biogenesis as a therapy for atherosclerosis. Nat Commun. 2017;8:15750. doi: 10.1038/ncomms15750

27. Ouimet M, Ediriweera HN, Gundra UM, Sheedy FJ, Ramkhelawon B, Hutchison SB, Rinehold K, van Solingen C, Fullerton MD, Cecchini K, et al. MicroRNA-33-dependent regulation of macrophage metabolism directs immune cell polarization in atherosclerosis. J Clin Invest. 2015;125:4334–4348. doi: 10.1172/JCI81676

28. Razani B, Feng C, Coleman T, Emanuel R, Wen H, Hwang S, Ting JP, Virgin HW, Kastan MB, Semenkovich CF. Autophagy links inflammasomes to atherosclerotic progression. Cell Metab. 2012;15:534–544. doi: 10.1016/j.cmet.2012.02.011

29. Tabas I. Apoptosis and efferocytosis in mouse models of atherosclerosis. Curr Drug Targets. 2007;8:1288–1296. doi: 10.2174/138945007783220623

30. Wang W, Liu W, Fidler T, Wang Y, Tang Y, Woods B, Welch C, Cai B, Silvestre-Roig C, Ai D, et al. Macrophage Inflammation, Erythrophagocytosis, and Accelerated Atherosclerosis in Jak2 (V617F) Mice. Circ Res. 2018;123:e35–e47. doi: 10.1161/CIRCRESAHA.118.313283

31. Yvan-Charvet L, Pagler TA, Seimon TA, Thorp E, Welch CL, Witztum JL, Tabas I, Tall AR. ABCA1 and ABCG1 protect against oxidative stress-induced macrophage apoptosis during efferocytosis. Circ Res. 2010;106:1861–1869. doi: 10.1161/CIRCRESAHA.110.217281

32. Kojima Y, Weissman IL, Leeper NJ. The Role of Efferocytosis in Atherosclerosis. Circulation. 2017;135:476–489. doi: 10.1161/CIRCULATIONAHA.116.025684

33. Jonsson AL, Backhed F. Role of gut microbiota in atherosclerosis. Nat Rev Cardiol. 2017;14:79–87. doi: 10.1038/nrcardio.2016.183

34. Wang Z, Klipfell E, Bennett BJ, Koeth R, Levison BS, Dugar B, Feldstein AE, Britt EB, Fu X, Chung YM, et al. Gut flora metabolism of phosphatidylcholine promotes cardiovascular disease. Nature. 2011;472:57–63. doi: 10.1038/nature09922

35. Warrier M, Shih DM, Burrows AC, Ferguson D, Gromovsky AD, Brown AL, Marshall S, McDaniel A, Schugar RC, Wang Z, et al. The TMAO-Generating Enzyme Flavin Monooxygenase 3 Is a Central Regulator of Cholesterol Balance. Cell Rep. 2015;10:326–338. doi: 10.1016/j.celrep.2014.12.036

36. Koeth RA, Wang Z, Levison BS, Buffa JA, Org E, Sheehy BT, Britt EB, Fu X, Wu Y, Li L, et al. Intestinal microbiota metabolism of L-carnitine, a nutrient in red meat, promotes atherosclerosis. Nat Med. 2013;19:576–585. doi: 10.1038/nm.3145

37. Rauch I, Deets KA, Ji DX, von Moltke J, Tenthorey JL, Lee AY, Philip NH, Ayres JS, Brodsky IE, Gronert K, Vance RE. NAIP-NLRC4 Inflammasomes Coordinate Intestinal Epithelial Cell Expulsion with Eicosanoid and IL-18 Release via Activation of Caspase-1 and −8. Immunity. 2017;46:649–659. doi: 10.1016/j.immuni.2017.03.016

38. Baetta R, Silva F, Comparato C, Uzzo M, Eberini I, Bellosta S, Donetti E, Corsini A. Perivascular carotid collar placement induces neointima formation and outward arterial remodeling in mice independent of apolipoprotein E deficiency or Western-type diet feeding. Atherosclerosis. 2007;195:e112–124. doi: 10.1016/j.atherosclerosis.2007.03.035

39. Dansky HM, Charlton SA, Sikes JL, Heath SC, Simantov R, Levin LF, Shu P, Moore KJ, Breslow JL, Smith JD. Genetic background determines the extent of atherosclerosis in ApoE-deficient mice. Arterioscler Thromb Vasc Biol. 1999;19:1960–1968. doi: 10.1161/01.atv.19.8.1960

40. de Villiers WJ, Smith JD, Miyata M, Dansky HM, Darley E, Gordon S. Macrophage phenotype in mice deficient in both macrophage-colony-stimulating factor (op) and apolipoprotein E. Arterioscler Thromb Vasc Biol. 1998;18:631–640. doi: 10.1161/01.atv.18.4.631

41. Bernier M, Mitchell SJ, Wahl D, Diaz A, Singh A, Seo W, Wang M, Ali A, Kaiser T, Price NL, et al. Disulfiram Treatment Normalizes Body Weight in Obese Mice. Cell Metab. 2020;32:203–214 e204. doi: 10.1016/j.cmet.2020.04.019

42. Baglione J, Smith JD. Quantitative assay for mouse atherosclerosis in the aortic root. Methods Mol Med. 2006;129:83–95. doi: 10.1385/1-59745-213-0:83

43. Zhao Y, Mahajan G, Kothapalli CR, Sun XL. Sialylation status and mechanical properties of THP-1 macrophages upon LPS stimulation. Biochem Biophys Res Commun. 2019;518:573–578. doi: 10.1016/j.bbrc.2019.08.089

44. Guz N, Dokukin M, Kalaparthi V, Sokolov I. If cell mechanics can be described by elastic modulus: study of different models and probes used in indentation experiments. Biophys J. 2014;107:564–575. doi: 10.1016/j.bpj.2014.06.033

45. Grant CA, Twigg PC, Saeed RF, Lawson G, Falconer RA, Shnyder SD. The Effect of Polysialic Acid Expression on Glioma Cell Nano-mechanics. Bionanoscience. 2016;6:81–84. doi: 10.1007/s12668-016-0192-2

46. Pierce M, Stuart J, Pungor A, Dryden P, Hlady V. Adhesion Force Measurements Using an Atomic Force Microscope Upgraded with a Linear Position Sensitive Detector. Langmuir. 1994;10:3217–3221. doi: 10.1021/la00021a053

47. Barot SV, Sangwan, N., Nair, K.G., Schmit, S., Xiang, S., Kamath, S.D., Liska, D., and Khorana, A.A. Tumor microbiome variation in young versus average onset colorectal cancer. Journal of Clinical Oncology. 2022;40:144–144. doi: 10.1200/JCO.2022.40.4_suppl.144

48. Chambers LM, Esakov Rhoades EL, Bharti R, Braley C, Tewari S, Trestan L, Alali Z, Bayik D, Lathia JD, Sangwan N, et al. Disruption of the Gut Microbiota Confers Cisplatin Resistance in Epithelial Ovarian Cancer. Cancer Res. 2022;82:4654–4669. doi: 10.1158/0008-5472.CAN-22-0455

49. Weiss K, Wanner N, Queisser K, Frimel M, Nunn T, Myshrall T, Sangwan N, Erzurum S, Asosingh K. Barrier Housing and Gender Effects on Allergic Airway Disease in a Murine House Dust Mite Model. Immunohorizons. 2021;5:33–47. doi: 10.4049/immunohorizons.2000096

50. Bolyen E, Rideout JR, Dillon MR, Bokulich NA, Abnet CC, Al-Ghalith GA, Alexander H, Alm EJ, Arumugam M, Asnicar F, et al. Reproducible, interactive, scalable and extensible microbiome data science using QIIME 2. Nat Biotechnol. 2019;37:852–857. doi: 10.1038/s41587-019-0209-9

51. Callahan BJ, McMurdie PJ, Rosen MJ, Han AW, Johnson AJ, Holmes SP. DADA2: High-resolution sample inference from Illumina amplicon data. Nat Methods. 2016;13:581–583. doi: 10.1038/nmeth.3869

52. McMurdie PJ, Holmes S. phyloseq: an R package for reproducible interactive analysis and graphics of microbiome census data. PLoS One. 2013;8:e61217. doi: 10.1371/journal.pone.0061217

53. Benjamini Y, and Hochberg, Y. Controlling the False Discovery Rate: A Practical and Powerful Approach to Multiple Testing. Journal of the Royal Statistical Society. . Journal of the Royal Statistical Society: Series B (Methodological). 1995;57:289–300.

54. Team RC. R: A language and environment for statistical computing. R Foundation for Statistical Computing. https://www.R-project.org/. 2018.

55. Hu JJ, Liu X, Xia S, Zhang Z, Zhang Y, Zhao J, Ruan J, Luo X, Lou X, Bai Y, et al. FDA-approved disulfiram inhibits pyroptosis by blocking gasdermin D pore formation. Nat Immunol. 2020;21:736–745. doi: 10.1038/s41590-020-0669-6

56. Martinez-Lopez N, Singh R. Autophagy and Lipid Droplets in the Liver. Annu Rev Nutr. 2015;35:215–237. doi: 10.1146/annurev-nutr-071813-105336

57. Filali-Mouncef Y, Hunter C, Roccio F, Zagkou S, Dupont N, Primard C, Proikas-Cezanne T, Reggiori F. The menage a trois of autophagy, lipid droplets and liver disease. Autophagy. 2022;18:50–72. doi: 10.1080/15548627.2021.1895658

58. Schrijvers DM, De Meyer GR, Kockx MM, Herman AG, Martinet W. Phagocytosis of apoptotic cells by macrophages is impaired in atherosclerosis. Arterioscler Thromb Vasc Biol. 2005;25:1256–1261. doi: 10.1161/01.ATV.0000166517.18801.a7

59. El-Kirat-Chatel S, Dufrene YF. Nanoscale adhesion forces between the fungal pathogen Candida albicans and macrophages. Nanoscale Horiz. 2016;1:69–74. doi: 10.1039/c5nh00049a

60. Patel NR, Bole M, Chen C, Hardin CC, Kho AT, Mih J, Deng L, Butler J, Tschumperlin D, Fredberg JJ, et al. Cell elasticity determines macrophage function. PLoS One. 2012;7:e41024. doi: 10.1371/journal.pone.0041024

61. Zhu W, Wang Z, Tang WHW, Hazen SL. Gut Microbe-Generated Trimethylamine N-Oxide From Dietary Choline Is Prothrombotic in Subjects. Circulation. 2017;135:1671–1673. doi: 10.1161/CIRCULATIONAHA.116.025338

62. Traughber CA, Iacano AJ, Neupane K, Khan MR, Opoku E, Nunn T, Prince A, Sangwan N, Hazen SL, Smith JD, Gulshan K. Impavido attenuates inflammation, reduces atherosclerosis, and alters gut microbiota in hyperlipidemic mice. iScience. 2023;26:106453. doi: 10.1016/j.isci.2023.106453

63. Frazier KR, Moore JA, Long TE. Antibacterial activity of disulfiram and its metabolites. J Appl Microbiol. 2019;126:79–86. doi: 10.1111/jam.14094

64. Shah O’Brien P, Xi Y, Miller JR, Brownell AL, Zeng Q, Yoo GH, Garshott DM, O’Brien MB, Galinato AE, Cai P, et al. Disulfiram (Antabuse) Activates ROS-Dependent ER Stress and Apoptosis in Oral Cavity Squamous Cell Carcinoma. J Clin Med. 2019;8. doi: 10.3390/jcm8050611

65. Rohatgi A. Reverse Cholesterol Transport and Atherosclerosis. Arterioscler Thromb Vasc Biol. 2019;39:2–4. doi: 10.1161/ATVBAHA.118.311978

66. Tall AR. Role of ABCA1 in cellular cholesterol efflux and reverse cholesterol transport. Arterioscler Thromb Vasc Biol. 2003;23:710–711. doi: 10.1161/01.ATV.0000068683.51375.59

67. Ouimet M, Barrett TJ, Fisher EA. HDL and Reverse Cholesterol Transport. Circ Res. 2019;124:1505-1518. doi: 10.1161/CIRCRESAHA.119.312617

68. Li J, Lin S, Vanhoutte PM, Woo CW, Xu A. Akkermansia Muciniphila Protects Against Atherosclerosis by Preventing Metabolic Endotoxemia-Induced Inflammation in Apoe-/- Mice. Circulation. 2016;133:2434–2446. doi: 10.1161/CIRCULATIONAHA.115.019645

69. Anand PK, Malireddi RK, Kanneganti TD. Role of the nlrp3 inflammasome in microbial infection. Front Microbiol. 2011;2:12. doi: 10.3389/fmicb.2011.00012

70. Puylaert P, Zurek M, Rayner KJ, De Meyer GRY, Martinet W. Regulated Necrosis in Atherosclerosis. Arterioscler Thromb Vasc Biol. 2022;42:1283–1306. doi: 10.1161/ATVBAHA.122.318177

71. Zhang J, Yu Q, Jiang D, Yu K, Yu W, Chi Z, Chen S, Li M, Yang D, Wang Z, et al. Epithelial Gasdermin D shapes the host-microbial interface by driving mucus layer formation. Sci Immunol. 2022;7:eabk2092. doi: 10.1126/sciimmunol.abk2092

72. Gao H, Cao M, Yao Y, Hu W, Sun H, Zhang Y, Zeng C, Tang J, Luan S, Chen P. Dysregulated Microbiota-Driven Gasdermin D Activation Promotes Colitis Development by Mediating IL-18 Release. Front Immunol. 2021;12:750841. doi: 10.3389/fimmu.2021.750841

73. Yan N, Wang L, Li Y, Wang T, Yang L, Yan R, Wang H, Jia S. Metformin intervention ameliorates AS in ApoE-/- mice through restoring gut dysbiosis and anti-inflammation. PLoS One. 2021;16:e0254321. doi: 10.1371/journal.pone.0254321

74. Chen ML, Yi L, Zhang Y, Zhou X, Ran L, Yang J, Zhu JD, Zhang QY, Mi MT. Resveratrol Attenuates Trimethylamine-N-Oxide (TMAO)-Induced Atherosclerosis by Regulating TMAO Synthesis and Bile Acid Metabolism via Remodeling of the Gut Microbiota. mBio. 2016;7:e02210–02215. doi: 10.1128/mBio.02210-15

75. Dai Z, Li S, Meng Y, Zhao Q, Zhang Y, Suonan Z, Sun Y, Shen Q, Liao X, Xue Y. Capsaicin Ameliorates High-Fat Diet-Induced Atherosclerosis in ApoE(-/-) Mice via Remodeling Gut Microbiota. Nutrients. 2022;14. doi: 10.3390/nu14204334

76. Rocha VZ, Libby P. Obesity, inflammation, and atherosclerosis. Nat Rev Cardiol. 2009;6:399-409. doi: 10.1038/nrcardio.2009.55

77. Fantuzzi G, Mazzone T. Adipose tissue and atherosclerosis: exploring the connection. Arterioscler Thromb Vasc Biol. 2007;27:996–1003. doi: 10.1161/ATVBAHA.106.131755

