## Supplementary material and methods for "Disulfiram reduces atherosclerosis and enhances efferocytosis, autophagy, and atheroprotective gut microbiota in hyperlipidemic mice"

Running title: DSF reduces atherosclerosis in hyperlipidemic mice.

C. Alicia Traughber[1,2,3], Kara Timinski[1,2], Ashutosh Prince[1,2], Nilam Bhandari[1,2], Kalash Neupane[1,2], Mariam R Khan[1,2], Esther Opoku[2], Emmanuel Opoku[3], Gregory Brubaker[3], Kanuri Nageshwar[4], Elif G. Ertugral[5], Prabha Naggareddy[4], Chandrasekhar R. Kothapalli[5], Jonathan D. Smith[3], and Kailash Gulshan[1,2,3]#

*Supplementary material and methods*

### Cell lines

Jurkat cells were maintained in DMEM (Cleveland Clinic Media Core #11-500p) supplemented with 10% FBS (Gibco #16140-071), 1% Pen/Strep 5000u/mL (Cleveland Clinic Media Core #721-100p). RAW-ASC (Invivogen #raw-asc) and RAW-ASC-KO-GSDMD (Invivogen #raw-kogsdmd) were maintained in DMEM (Cleveland Clinic Media Core #11-500p) supplemented with 10% FBS, 1% Pen/Strep 5000u/mL, 100 μg/ml Blasticidin (Invivogen #ant-bl) and 100 μg/ml Normocin (Invivogen #ant-nr) RAW-Difluo™ mLC3 (Invivogen # awdf-mlc3) were maintained in DMEM supplemented with 10% FBS (Gibco), 1% pen-strep, 100 μg/ml Zeocin (Invivogen #ant-zn) and 100 μg/ml Normocin (Invivogen). THP-1 cells (ATCC # TIB 202) were maintained in RPMI 1640 (Cleveland Clinic Media Core #10-500p) containing 10% FBS, 1% Pen/Strep, 0.05 mM 2-mercaptoethanol (Sigma #M3148). THP-1 cells were differentiated into macrophages (THP-1 macrophages) using 100ng/mL phorbol 12-myristate 13-acetate (Sigma P8139) for 3 days. HepG2 cells (ATCC) were maintained in EMEM (Cleveland Clinic Media Core #99BJ500CUSTp) supplemented with 10% FBS and 1% Pen/Strep 5000u/ml. All cells were maintained at 37°C with 5% $CO_2$.

### Isolation and maturation of bone-marrow derived macrophages

C57BL/6J-WT or C57BL/6J-GsdmD$^{-/-}$ mice were maintained on standard chow diet and water. Mice were euthanized by $CO_2$ inhalation and femoral bones were removed. The marrow was flushed out of the bones into a 50 ml sterile tube using a 10 ml syringe with a 26-gauge needle filled with sterile DMEM. Cells were centrifuged for 5 min at 1,800 rpm at 4°C, followed by two washes with sterile PBS. The cells were resuspended in sterile-filtered BMDM growth media (DMEM with 7.6% fetal bovine serum, 15% L-cell conditioned media, and 0.76% penicillin/streptomycin mixture) and plated in 10 cm culture dishes and incubated at 37°C for 14 days, with media changes every 2-3 days.

**In-vivo and in vitro inflammasome assembly**

For in-vivo inflammasome study, 4–5-week-old mice were fed a WTD or DSF diet for three weeks, and i.p injected with sterile 5μg LPS (MilliporeSigma) for 4h. The Nlrp3 inflammasome assembly in mice was induced by i.p injection of ATP (0.5 ml of 30 mM, pH 7.0). The mice were euthanized after 30 min and peritoneal cavity was lavaged with 5 ml PBS. Approximately 3.5 ml peritoneal lavage fluid was recovered from each mouse and centrifuged at 15K rpm for 10 min at room temperature. The supernatant was subjected to IL-1β ELISA, using mouse IL-1β Quantikine ELISA kit (R&D systems # MLB00C,) following manufacturer's instructions. For in-vitro Nlrp3 activity, cells were plated at equal densities and primed with 1 µg/ml LPS (MilliporeSigma #L2880) for 3-4 hours and/or pretreated ± 5 and 10µM Disulfiram (Sigma #PHR1690) and stimulated with 1 mM ATP (MilliporeSigma #A2383) for 30m or in serum free media. Media was harvested and subjected to IL-1β ELISA according to manufacturer's instructions (R&D Systems #MLB00C)

**Viability Assay**

Cells were plated in equal number in 24-well format. Cells were treated with± 2, 5, or 10µM DSF or 0.5% digitonin for 2, 6, and 24 hours. At end of each time point, cells were treated with Live/dead fixable blue stain, then wash twice with PBS, then viability was quantified using iD3 SpectraMax microplate reader (Molecular devices), excited at 350nm and emission recorded at 450nm.

**Cytotoxicity Assay**

Cells were plated at 10,000 cells / well in a 96-well format. Cells were treated with ± 2, 5, or 10µM DSF for 0, 6, and 24 hours. Spontaneous LDH Activity Control (S-LDH) and

Maximum LDH Activity Control (M-LDH) were prepared according to manufacturer's instructions. Media after each time point was collected at used in the CyQuant LDH Cytotoxicity Assay according to manufacturer's instructions (Thermofisher cat# C20300). Absorbance for cytotoxicity was measured using the iD3 SpectraMax microplate reader (Molecular devices), Absorbance = 490nm absorbance – 680nm absorbance.

**Western Blotting**

Western blot analysis was performed on protein extracts ± various treatment conditions. Equal amounts of proteins from tissue or cells, determine by either BCA assay (Pierce) Nanodrop 2000 (Thermo Scientific), were resolved on a Novex 4-20% Tris-Glycine Gel (Invitrogen) then transferred to a PVDF membrane (Invitrogen). After blocking in Casein (Thermo Scientific #37525), the blots were probed with respective primary antibodies 1:1000 for overnight, washed with PBS + Tween 20 (Fisher Chemical #BP337-500) then probed with a 1:10,000 dilution of respective secondary antibody for 1 hour, then washed with PBST. Blots were developed using the Immobilon Western Chemiluminescent HRP Substrate (Millipore #WBKLS05000 and imaged using the iBright™ CL750 Imaging System (Invitrogen #A44116).

**Autophagy Reporter Assay**

RAW-Difluo mLC3 were plated in chamber slides (Ibidi; #80427). Cells were treated with ± 5 μM DSF for 2h then washed with 3x with PBS, fixed in 3.7% paraformaldehyde for 3m, washed with PBS, and then counterstained for DAPI (Invitrogen #S36964) and mounted. Microscopy was performed using the Nikon Eclipse Ti confocal and Nikon NIS Elements Imaging Software version 4.1.

**Indirect Immunofluorescence**

THP-1 macrophages were treated with ± 5 μM DSF for 2h, then fixed with 4% paraformaldehyde. Cells were blocked with 5% goat serum and 1% BSA in PBS. Cells were incubated with 1:25 dilution of MerTK primary antibody (Thermo PA5-15028) for overnight, washed with PBS, then incubated with 1:400 dilution goat-anti-rabbit AlexaFluor488 for 1h, washed with PBS, then stained for DAPI (Molecular Probes #36964) and imaged using the Nikon Eclipse T*i* confocal and Nikon NIS Elements Imaging Software version 4.13.

**Antibody table**

| Antibodies | Dilution | Vendor |
|---|---|---|
| LC3A/B (D3U4C) XP® Rabbit mAb | 1:50 | Cell Signaling Technology #12741 |
| LC3B (E5Q2K) Mouse mAb #83506 | 1:1000 | Cell Signaling Technology #83506 |
| p62/SQSTM1 Antibody - BSA Free | 1:1000 | Novus Biologicals #NBP1-48320 |
| Goat anti-Rabbit IgG (H+L) Highly Cross-Adsorbed Secondary Antibody, Alexa Fluor™ 488 | 1:150 | ThermoFisher Scientific #A11034 |
| Goat anti-Rabbit IgG (H+L) Cross-Adsorbed Secondary Antibody, Alexa Fluor™ 488 | 1:400 | ThermoFisher Scientific #A11008 |
| β-Actin (C4) | 1:5000 | Santa Cruz Biotech #sc-47778 |
| Goat Anti-Mouse IgG Antibody, (H+L) HRP conjugate | 1:10,000 | MilliporeSigma #AP308P |
| Goat Anti-Rabbit IgG Antibody, (H+L) HRP conjugate | 1:10,000 | MilliporeSigma #AP307P |
| ASC/TMS1 (D2W8U) Rabbit mAb | 1:1000 | Cell Signaling Technology; #67824 |
| MerTK Polyclonal Antibody | 1:25 | ThermoFisher Scientific PA5-15028 |

**Supplemental Figures and Legends**

***Fig. S1***

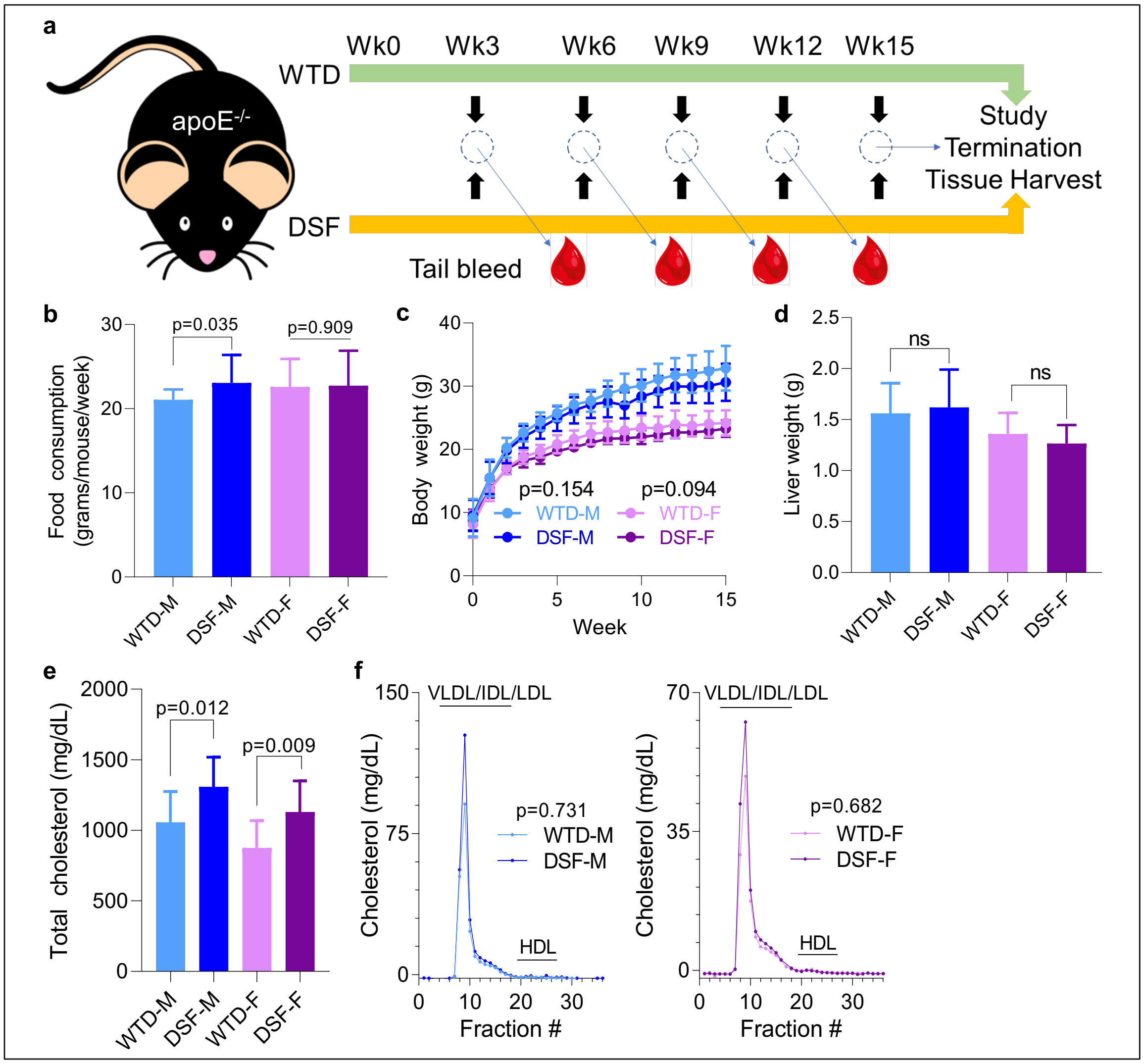


**Fig. S1. a)** Schematic of DSF study showing that apoE$^{-/-}$ mice were weaned onto WTD ± DSF for 15 weeks; n=9-13 mice / group and sex**. b)** Food consumption in mice fed with WTD ± DSF (p-values are determined by t-test). **c)** Weekly body weight of mice fed with WTD ± DSF (p-values are determined by 2-way ANOVA). **d)** Male (M) and female (F) liver weight from 15-week WTD± DSF-fed mice, p-value determined by t-test. **e)** Total plasma cholesterol levels in mice fed with WTD ± DSF (p- values are determined by t-test). **f)** FPLC showing cholesterol peaks of pooled plasma from males and females fed with WTD ± DSF for 15 weeks (p-values are determined by t-test).

### Fig. S2

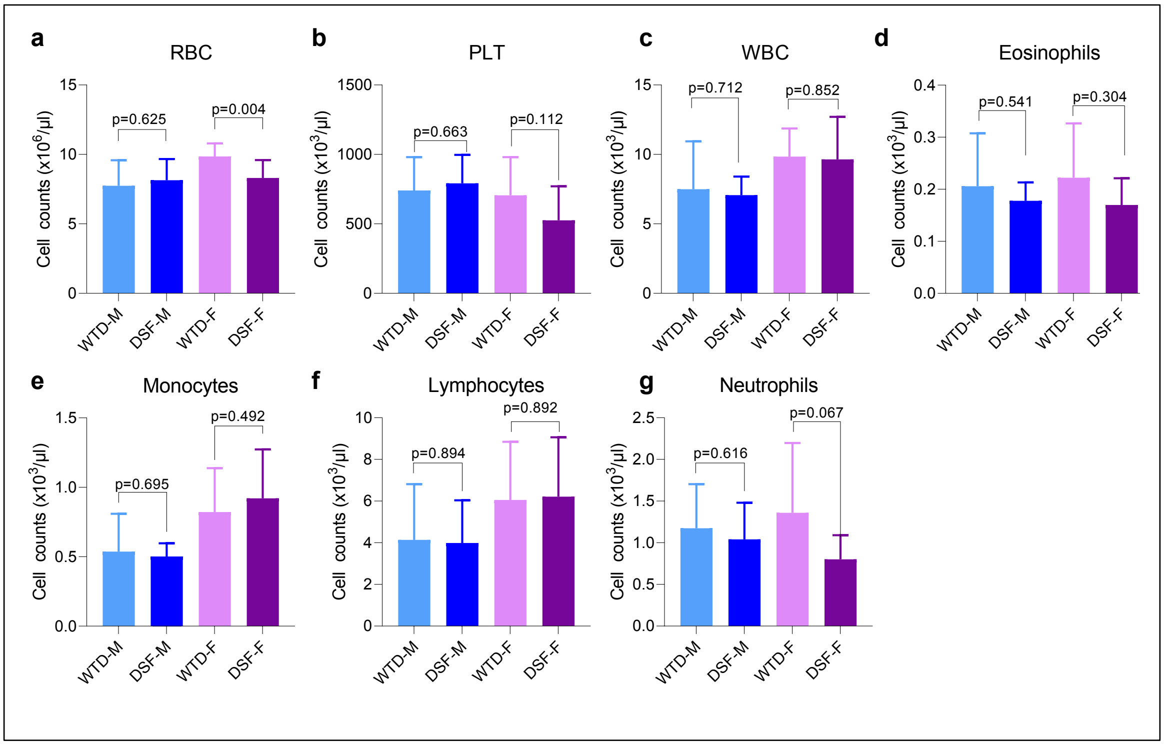


**Fig. S2. DSF effects on complete blood counts (CBC) in hyperlipidemic apoE$^{-/-}$ mice.** Male (M) and female (F) CBC from 15-week WTD± DSF-fed mice; n=7-12 mice / group and sex, p-values determined by t-test. **a)** Red blood cells (RBC). **b)** Platelets (PLT). **c)** White blood cells (WBC). **d)** Eosinophils. **e)** Monocytes. **f)** Lymphocytes. **g)** Neutrophils.

### Fig. S3

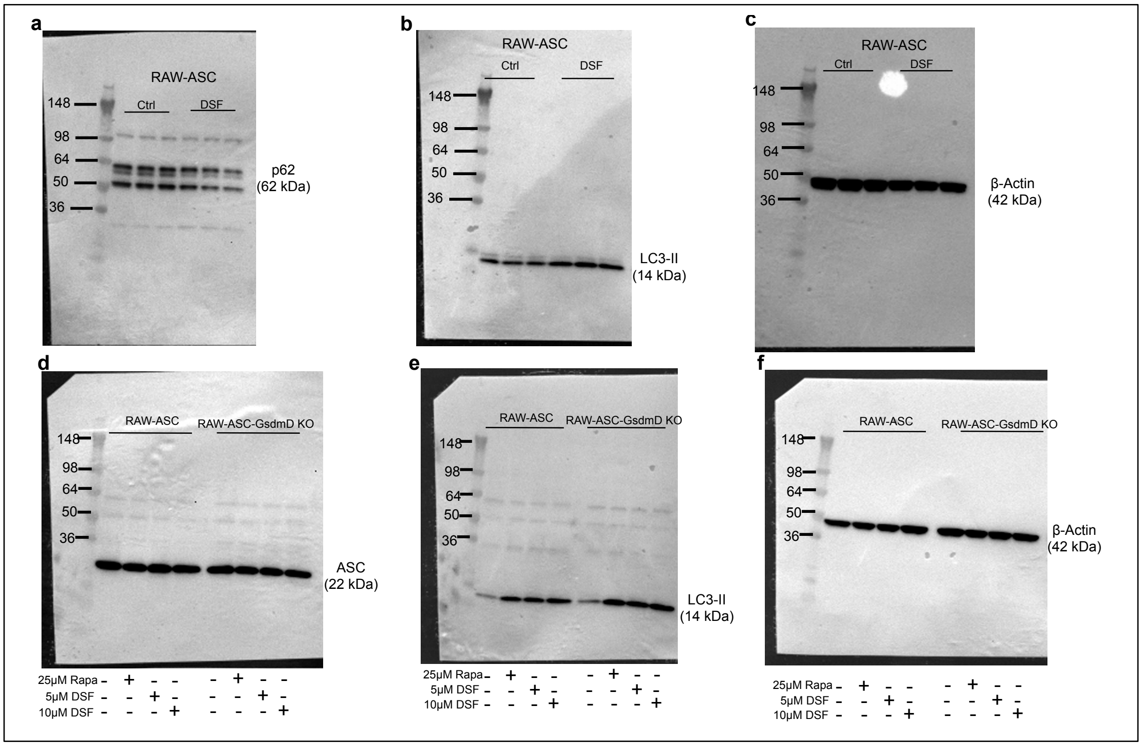


**Fig. S3. Original Blots for main Figure 4.** RAW-ASC macrophages were treated with ±5 μM DSF for 2h; n=3. **a**) Western blot of p62 **b**) Relative expression of LC3-II. **c**) Western blot of β-actin **d**) Western blot of ASC in macrophages treated with ± Rapamcyin or ± DSF for 2h **e**) Western blot of LC3-II. **f**) Western blot of β-actin.

**Fig. S4**

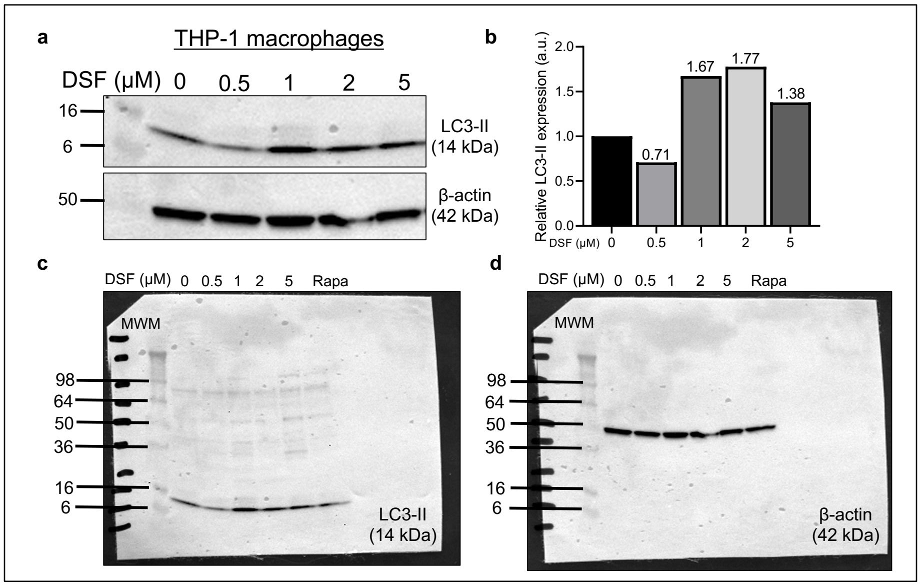


**Fig. S4. DSF-induced autophagy in THP-1 macrophages.** THP-1 macrophages were treated with 0, 0.5, 1, 2, 5 μM DSF for 2h. **a**) Western blot of LC3-II expression. **b**) Relative expression of LC3-II vs. β-actin. **c, d**) Original Western blots of ***S4a***.

**Fig. S5**

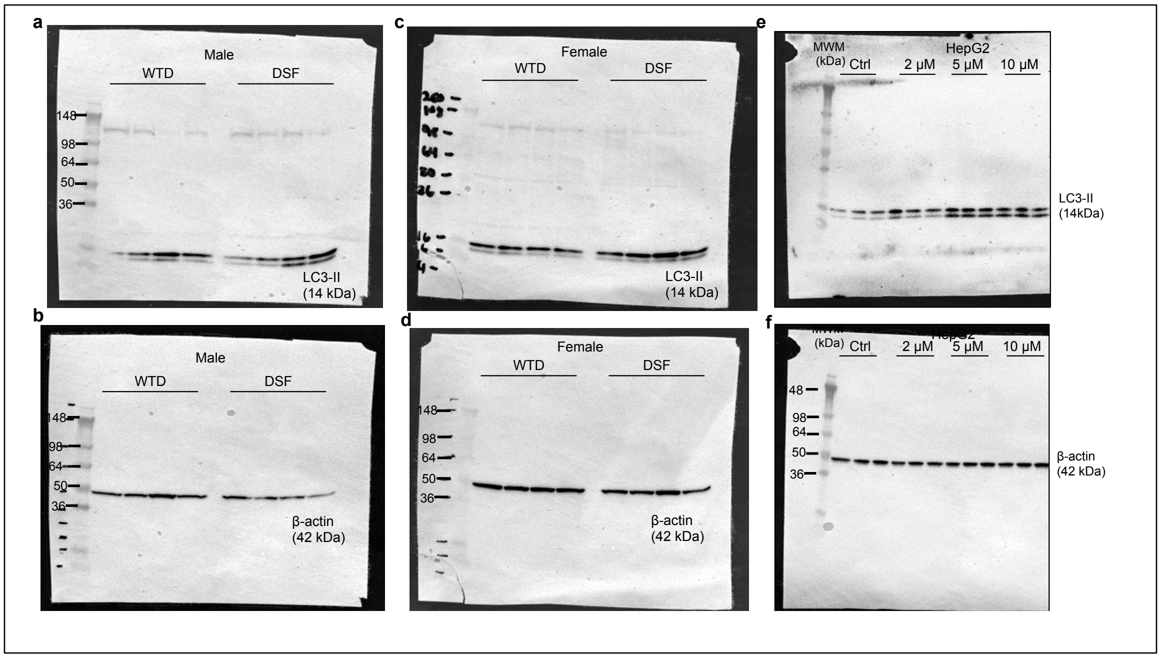


**Fig. S5. Original Western Blots for main Figure 5. a,b**) Male and **c,d**) female liver western blot of LC3-II expression n=4/group and sex. Western blot of HepG2 cells ± DSF (2,5,10 μM, n=3) showing LC3-II (***e***) and β-actin (**f**).

**Fig. S6**

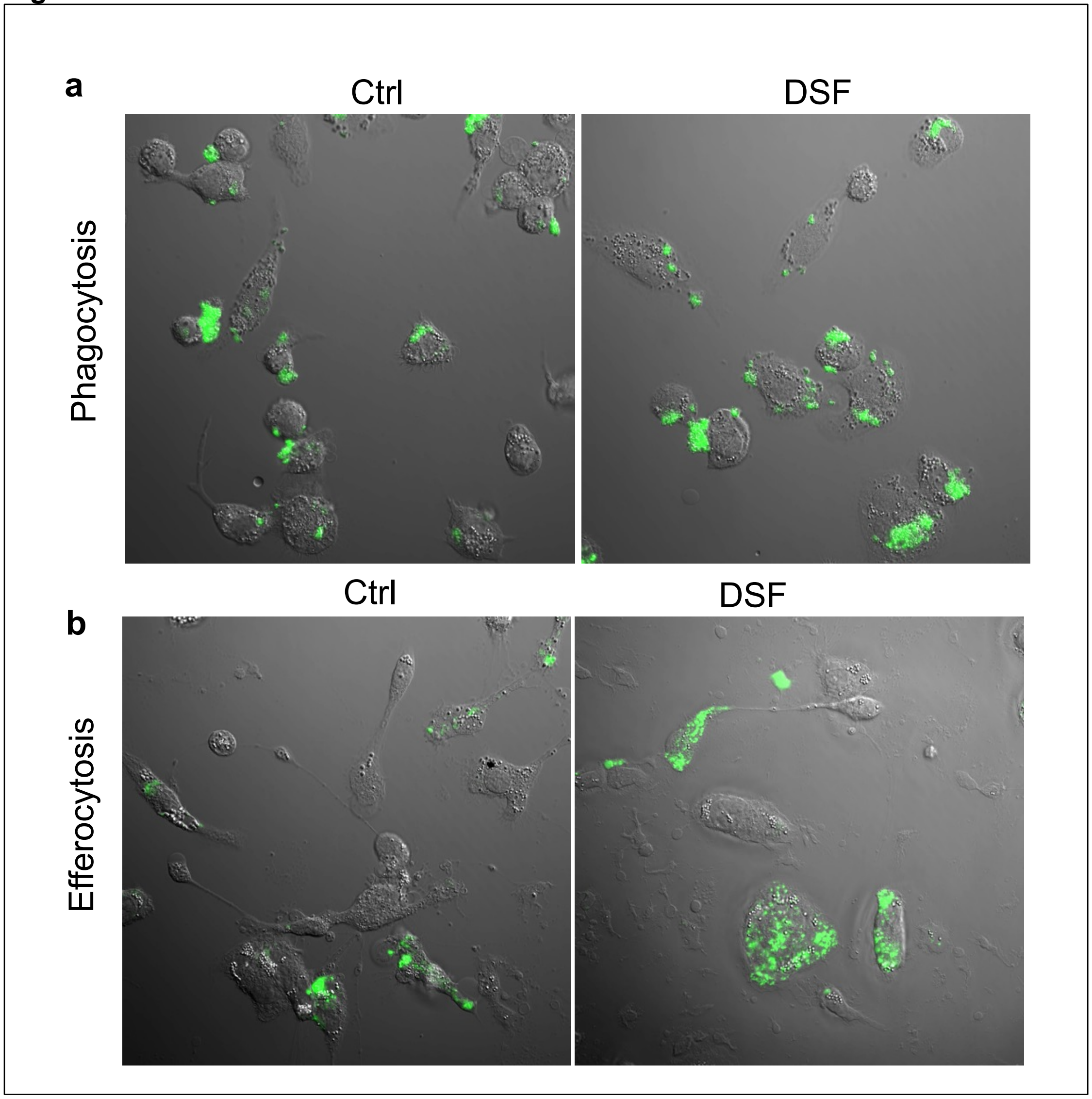


**Fig. S6. Original Field of cells for main Figure 5, a)** Phagocytosis (Original figure 5a), **b**) Efferocytosis (Original figure 5c).
